# CALX-CBD1 Ca^2+^-binding cooperativity studied by NMR spectroscopy and ITC with Bayesian statistics

**DOI:** 10.1101/791178

**Authors:** M. V. C. Cardoso, J. D. Rivera, P. M. Vitale, M. F. S. Degenhardt, L.A. Abiko, C. L. P. Oliveira, R. K. Salinas

## Abstract

The Na^+^/Ca^2+^ exchanger of *Drosophila melanogaster*, CALX, is the main Ca^2+^-extrusion mechanism in olfactory sensory neurons and photoreceptor cells. Na^+^/Ca^2+^ exchangers have two Ca^2+^ sensor domains, CBD1 and CBD2. In contrast to the mammalian homologues, CALX is inhibited by Ca^2+^-binding to CALX-CBD1, while CALX-CBD2 does not bind Ca^2+^ at physiological concentrations. CALX-CBD1 consists of a *β*-sandwich and displays four Ca^2+^ binding sites at the tip of the domain. In this study, we used NMR spectroscopy and isothermal titration calorimetry (ITC) to investigate the cooperativity of Ca^2+^-binding to CALX-CBD1. We observed that this domain binds Ca^2+^ in the slow exchange regime at the NMR chemical shift time scale. Ca^2+^-binding restricts the dynamics in the Ca^2+^-binding region. Experiments of ^15^N CEST and ^15^N R_2_ dispersion allowed the determination of Ca^2+^ dissociation rates (≈ 20 s^−1^). NMR titration curves of residues in the Ca^2+^-binding region were sigmoidal due to the contribution of chemical exchange to transverse magnetization relaxation rates, R_2_. Hence, a novel approach to analyze NMR titration curves was proposed. Ca^2+^-binding cooperativity was examined assuming two different stoichiometric binding models and using a Bayesian approach for data analysis. Fittings of NMR and ITC binding curves to the Hill model yielded *n*_Hill_ = 2.9 − 3.1, near maximum cooperativity (*n*_Hill_ = 4). By assuming a stepwise model to interpret the ITC data, we found that the probability of binding from 2 up to 4 Ca^2+^ is at least three orders of magnitude higher than that of binding a single Ca^2+^. Hence, four Ca^2+^ ions bind almost simultaneously to CALX-CBD1. Cooperative Ca^2+^-binding is key to enable this exchanger to efficiently respond to changes in the intracellular Ca^2+^-concentration in sensory neuronal cells.

**SIGNIFICANCE:** CALX-CBD1 is the Ca^2+^-sensor domain of the Na^+^/Ca^2+^ exchanger of *Drosophila melanogaster*. It consists of a *β*-sandwich, and contains four Ca^2+^ binding sites at the distal loops. In this study, we examined the cooperative binding of four Ca^2+^ ions to CALX-CBD1 using NMR spectroscopy and isothermal titration calorimetry (ITC) experiments. NMR and ITC data were analyzed using the framework of the binding polynomial formalism and Bayesian statistics. A novel approach to analyze NMR titration data in the slow exchange regime was proposed. These results support the view that CALX-CBD1 binds four Ca^2+^ with high cooperativity. The significant ligand binding cooperativity exhibited by this domain is determinant for the efficient allosteric regulation of this exchanger by intracellular Ca^2+^.

## INTRODUCTION

Ca^2+^ is a key intracellular regulator and an important second messanger (1). Hence, cells developed various mechanisms to control the intracellular Ca^2+^ concentration very precisely (2, 3). Among those, the Na^+^/Ca^2+^ exchanger (NCX) is particularly relevant (4). After the Ca^2+^-ATPases, the NCX is one of the most important mechanisms for the extrusion of intracellular Ca^2+^, particularly in excitable cells (5). The NCX couples the translocation of one Ca^2+^ to the counter-transport of three Na^+^ across the cell membrane (5, 6). It consists of a large transmembrane domain, involved in ion translocation across the lipid bilayer, and a large intracellular loop that connects transmembrane helices 5 and 6 and that is responsible for the regulation of exchanger activity. Binding of Ca^2+^ to two calcium-binding domains (CBD1 and CBD2), located at the NCX cytoplasmic loop, activates the exchanger, enabling the in/out flux of Na^+^ and Ca^2+^. The Na^+^/Ca^2+^ exchanger of *Drosophila melanogaster*, so-called CALX, is unusual, since Ca^2+^-binding to the CBDs inhibits this exchanger (7). This behavior is curious because the CALX and the NCX display approximately 44 % of amino acid sequence identity (8), and are believed to share the same architecture. The CALX Na^+^/Ca^2+^ exchanger is found in *Drosophila* photoreceptor cells and in olfatory sensory neurons, where it is the main Ca^2+^ extrusion mechanism (9, 10). CALX plays an essential role in light-mediated signalling. Mutation of CALX in transgenic flies caused a reduction in light signal response amplification, while its overexpression generated significant retinal degeneration (9).

The CBDs are also called CALX-*β* motifs (8) or CALX-*β* domains (11). They appear adjacent to each other, separated by a linker of three amino acids, which is highly conserved among exchangers of different organisms (12, 13). The CALX-CBD2 domain does not bind Ca^2+^ at physiological concentrations (14). Hence, in the CALX exchanger, the regulatory Ca^2+^-binding sites are located only in the CBD1 domain (13). The CALX-CBD1 crystal structure shows four Ca^2+^ in the distal loops of the CBD1 *β*-sandwich, at the tip of the domain (15) (Fig. 1A). The four Ca^2+^-binding sites are located at approximately 4Å from each other. A series of negatively charged residues provides most of the anchoring points for Ca^2+^ coordination. Most of these residues are located in the EF loop and at the C-terminus (Fig. 1A), near the linker with CBD2. CALX-CBD1 was also crystallized in the Apo state and in an intermediate state (15). While the Apo state structure shows no electron density for the EF loop, the crystal structure of the intermediate state shows two Ca^2+^ at sites Ca1 and Ca2. The intermediate state structure is very similar to the fully loaded Ca^2+^-bound state, with a backbone RMSD of only 0.3Å. These observations indicate that the binding of Ca^2+^ to sites Ca1 and Ca2 is sufficient to stabilize the structure of the Ca^2+^-binding region of CALX-CBD1 (15). Furthermore, they suggest that the Ca^2+^-binding mechanism is sequential, rather than the simultaneous binding of four Ca^2+^ to sites Ca1-Ca4 in all-or-none fashion (15).

**Figure 1:**
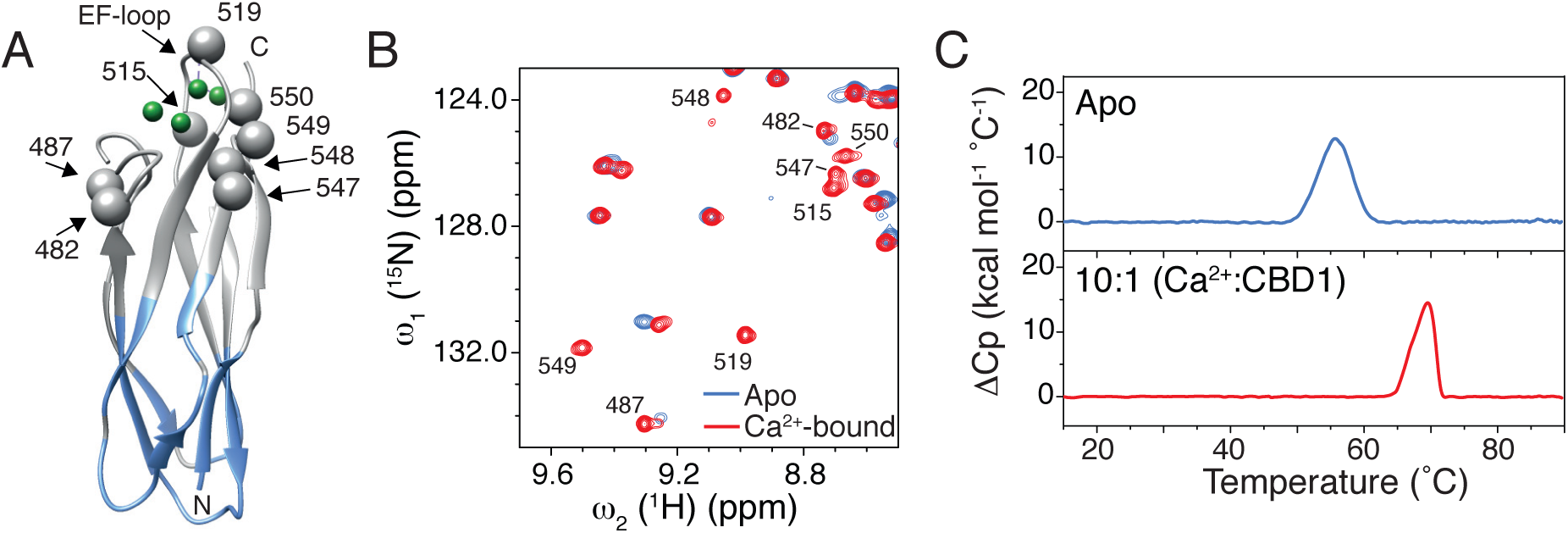
Binding of Ca^2+^ stabilizes CALX-CBD1. A) Crystal structure of CALX-CBD1 in the Holo state (PDB 3EAD (15)). Ca^2+^ ions are shown as green spheres. Residues whose ^1^H^15^N cross peaks appear only in the presence of Ca^2+^ are indicated and shown as gray spheres. The assigned residues in the unbound state are shown in blue. B) Superposition of ^1^H-^15^N HSQC spectra of CALX-CBD1 recorded in the absence (blue) and presence (red) of 20 mM of CaCl_2_. C) Differential scanning calorimetry experiment showing the thermal denaturation of CALX-CBD1 in the absence (blue), and presence of Ca^2+^ (red). The protein melting temperatures, T_m_, under these conditions are 55.3 °C and 69.6 °C for the Apo state and Ca^2+^-bound states, respectively.

The Ca^2+^-binding cooperativity exihibited by the CBDs is critical to enable them to function as efficient allosteric switches of the Na^+^/Ca^2+^ exchangers. Therefore, in this work, we used NMR spectroscopy and isothermal titration calorimetry (ITC) experiments to investigate the Ca^2+^-binding properties of CALX-CBD1. Consistent with the crystallographic data, we observed that CALX-CBD1 displays substantial backbone flexibility near the Ca^2+^-binding sites. Experiments of ^15^N chemical exchange saturation transfer (CEST) and of ^15^N Carr-Purcell-Meiboom-Gill (CPMG) relaxation dispersion, recorded at the Ca^2+^:CALX-CBD1 stoichiometric molar ratio, allowed the determination of Ca^2+^ dissociation rates. Ca^2+^-binding cooperativity was examined by isothermal titration calorimetry (ITC) and NMR titration experiments, assuming different stoichiometric binding models, and using a Bayesean approach for data analysis. The NMR titration curves obtained for residues near the Ca^2+^ binding sites displayed an accentuated sigmoidal behavior, which could be accounted for by calculating the effective transverse magnetization relaxation rates, 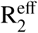, at each titration point, using the exact solution of the Bloch-McConnell equation (16). The analysis of NMR and ITC data converged to the view that CALX-CBD1 binds four Ca^2+^ ions in a highly cooperative fashion. Indeed, the probability of binding from 2 up to 4 Ca^2+^ is at least three orders of magnitude higher than that of binding a single Ca^2+^. In summary, this study sheds new light on the Ca^2+^-binding cooperativity of CALX-CBD1. Ligand binding cooperativity is an important characteristic of a biological switch. This characteristic enables the Na^+^/Ca^2+^ exchangers to efficiently respond to changes in the intracellular Ca^2+^ concentration, and, hence, to efficiently control intracelluslar Ca^2+^ homeostasis.

## NMR TITRATION THEORETICAL FRAMEWORK

In a NMR titration experiment, in which the protein-ligand binding equilibrium follows the slow exchange regime, a titration curve may be built by computing the variable η, the intensity ratio of the protein NMR signal in the bound state at a given ligand concentration, *I*, and at saturating conditions, *I*_max_:

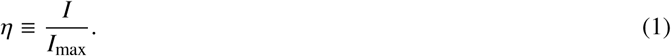

Assuming a Lorentzian lineshape, the maximum intensity of a ^1^H-^15^N HSQC cross peak may be written as:

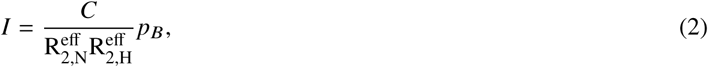

where *C* is a constant that takes into account the physical dimensions of the equilibrium magnetization, 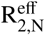 and 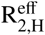 are the effective transverse relaxation rates for ^15^N and ^1^H nuclei, respectively, and *p*_*B*_ is the population of the protein molecules in the *bound* state. One may note that an analogous binding curve could be derived by monitoring the intensity decrease of the protein NMR signals in the Apo state.

Using the exact solution for R_2_ in chemically exchanging systems derived by Leigh (1971) (16), and assuming equivalent intrinsic 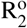 relaxation rates for the two exchanging states, we can write 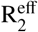 as:

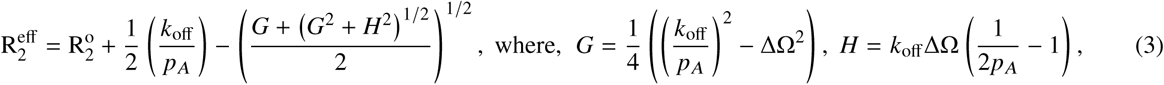

*p*_*A*_ is the population of protein in the Apo state, ΔΩ is the chemical shift difference between the bound and Apo states, and *k*_off_ is the ligand dissociation rate constant. Under conditions of saturation, i.e. when all ligand binding sites are occupied, 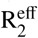 tends to 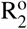, hence the exchange contribution to the effective transverse relaxation rate is negligible and the cross peak intensity is given by:

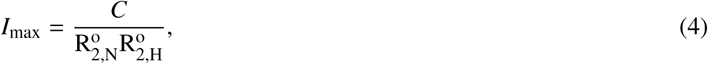

Therefore, it is convenient to rewrite the variable *η* as:

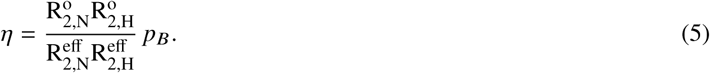

The populations of protein molecules in the Apo and in the bound states may be written in terms of the binding polynomial (Eq. S6 in the Supporting Material):

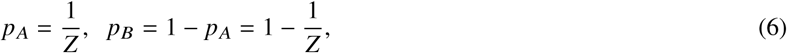

One should note that the population *p*_*B*_, as defined here, contains the contribution of all protein bound states. By substituting Eqs. 6 into Eq. 5 we may re-state *η* as:

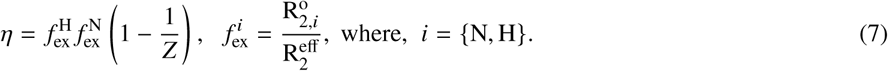

## METHODS

### CALX-CBD1 expression and purification

A DNA fragment corresponding to residues 434-555 from the *Drosophilia melanogaster* CALX 1.1, was PCR amplified from the *CALX1.1* cDNA, and cloned in fusion with a N-terminal six-histidines tag in the vector pET-28a (Novagen), using NdeI and XhoI restriction sites. Protein expression was carried out in *Escherichia coli* BL21-CodonPlus (DE3)-RIL (Agilent). Uniform labeling with ^15^N, or with ^15^N and ^13^C, was achieved by overexpression in M9 supplemented with 1.0 g/l of ^15^NH_4_Cl (Cambridge Isotopes Laboratories, CIL) and/or 4.0 g/l of ^13^C-labeled glucose (CIL). Bacterial cultures were grown at 37 °C to an OD_600_ of 0.7, at which point the temperature was decreased to 18 °C, and protein overexpression was induced by the addition of 0.4 mM of isopropyl-*β*-D-thiogalactopyranoside (IPTG) overnight. Cells were harvested by centrifugation, and the cell pellets stored at −20 °C. Approximately 10 g of cells were suspended in 100 ml of lysis buffer (20 mM Tris pH 7.0, 200 mM NaCl, 5 mM *β*-mercaptoetanol, 0.1 µg/ml of pepstatin and aprotinin, 1.0 mM PMSF, and 1 mg/ml lysozyme), and subsequently lysed by sonication using a VCX 500 (Sonics) instrument. The cell lysate was clarified by centrifugation at 21000 g for one hour, and the supernantant was applied into a 5 ml Ni^2+^ column (HisTrap, GE Healthcare) pre-equilibrated with buffer A (20 mM sodium phosphate pH 7.4, 300 mM NaCl, 5 mM *β*-mercaptoethanol, 10 mM Imidazole), washed with 5 column volumes of buffer A, and eluted with 0.5 M of Imidazole. Fractions containing CALX-CBD1 were combined, concentrated to 2 ml using an Amicon concentrator with a 3 kDa cutoff (Millipore), and applied into a gel filtration Superdex 75 (16/600) column (GE Healthcare) pre-equilibrated with the gel filtration buffer (20 mM Tris pH 8.0, 200 mM NaCl, 5 mM *β*-mercaptoethanol, 1% (V/V) glycerol). Finally, the desired fractions were combined, and applied into a MonoQ 10/100 (GE Healthcare) column, pre-equilibrated with the gel filtration buffer, and eluted with a gradient of 1.0 M of NaCl. In order to prepare NMR samples, the eluted CALX-CBD1 was concentrated and buffer exchanged to 20 mM Tris-HCl pH 7.4, containing 5 mM of *β*-mercaptoethanol, 200 mM of NaCl, 1% (V/V) of glycerol, and 10% of D_2_O. Protein concentration was determined by measuring the absorbance at 280 nm assuming *ϵ*_280_ = 7575 M^−1^cm^−1^. The molecular weight of CALX-CBD1 was confirmed as being 16.025 kDa using mass spectrometry (MALDI-TOF-TOF).

### NMR spectroscopy

All NMR spectra were recorded on a Bruker AVANCE III spectrometer equipped with a TCI cryoprobe, and operating at 800 MHz (^1^H frequency). CPMG-based ^15^N relaxation dispersion experiments were additionally recorded on a Bruker Avance III-HD spectrometer operating at 600 MHz (^1^H frequency) and a equipped with a TCI cryoprobe. NMR experiments were carried out at 308 K unless otherwise stated. All NMR data were processed with NMRPipe (17), and analyzed using the CcpNmr Analysis software (18). Protein samples used for backbone resonance assignment consisted of approximately 200 *µ*M of ^15^N/^13^C double-labeled CALX-CBD1. The Apo state sample contained 2 mM of EDTA, while the Ca^2+^-bound sample contained 20 mM of CaCl_2_. Backbone (^1^HN, ^15^N, ^13^C*α*, ^13^CO) and side-chain (^13^C*β*) resonance assignments for CALX-CBD1 were obtained from the analysis of 3D HNCA, HN(CO)CA, HNCO, HN(CA)CO, CBCA(CO)NH, and HNCACB experiments. The assigned chemical shifts for CALX-CBD1 in the Ca^2+^-bound state were deposited in the BMRB under the accession code 27787.

Longitudinal R_1_ and rotating-frame (R_1*ρ*_) relaxation rates, and the [^1^H]-^15^N heteronuclear NOE, were measured using standard ^15^N relaxation methods recorded as pseudo 3D or interleaved experiments (19, 20). Protein samples consisted of 380 *µ*M of ^15^N-labeled CALX-CBD1 containing CaCl_2_ at a molar ratio of 16:1 (CaCl_2_:protein). The R_1*ρ*_ experiment was recorded using spin-lock field strength γ*B*_1_ / 2π = 1013 Hz. Rotating frame R_1*ρ*_ relaxation was sampled at 4, 8, 16, 24, 35, 44, 58, 66, 80, 92, 104, 115 ms, while longitudinal R_1_ relaxation was sampled at 150, 300, 450, 600, 900, 1250, 1500, 1800, 2000, 2250 and 2500 ms. The heteronuclear NOE was recorded using a saturation time of 3.0 s. The recycle delay was set to 4.0 s for the R_1_ and R_1*ρ*_ experiments, while a delay of 5.0 s was used for the heteronuclear NOE experiment. R_1_ and R_1*ρ*_ relaxation rates were determined by fitting the cross peak intensity decays to an exponential decay function consisting of two parameters, the intensity at time zero and a decay rate. The uncertainties of the peak heights were taken from the noise level in each spectrum, while the uncertainties of the fitted parameters were estimated using Monte Carlo (at least 1000 iterations). R_1*ρ*_ relaxation rates were measured at two different ^15^N carrier frequencies, 123.942 and 110.942 ppm, satisfying the conditions presented by Massi et. al. (2004) (20). The chosen R_1*ρ*_ values corresponded to the smallest Ω / (*γB*_1_) ratio, in which Ω is the resonance offset in Hz, indicating best alignment of the magnetization with the spin-lock field. R_2_ values were obtained by correcting R_1*ρ*_ for off-resonance effects using the following relation: R_2_ = R_1*ρ*_ / sin^2^ *θ* − R_1_ (cos *θ* / sin *θ*)^2^, where *θ* = arctan (*γB*_1_ / Ω).

A relaxation-compensated Bruker pulse sequence (hsqcrexetf3gpsitc3d) was used for all CPMG-based ^15^N relaxation dispersion experiments, using constant relaxation times of 40 ms or 50 ms. The experiments were recorded as a series of 2D planes with CPMG field strenghts of 50, 100, 150, 200, 250, 300, 350, 400, 500, 600, 700, 800 and 900 Hz, and a reference spectrum measured by omitting the CPMG period. The cross-peak intensities were converted into effective transverse relaxation rates doing 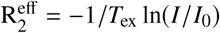, where *I*_0_ is the cross-peak intensity in the reference spectrum. Uncertainties of 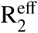 were calculated by error propagation assuming the noise level as the uncertainty in the cross-peak heights in each spectrum. CPMG relaxation dispersion curves recorded at 600 and 800 MHz were simultaneously fitted to the Bloch-McConnell equation assuming a two-site chemical exchange model, through the calculation of the following equation using a home-made python script:

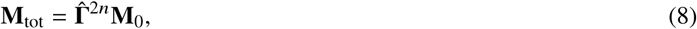

where **M**_0_ is the initial transverse magnetization aligned with the *x*-axis, *n* is the number of CPMG periods, and 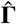 is given by:

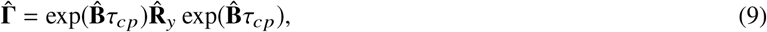

where 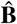 is the Bloch-McConnell equation for the transverse magnetization, and 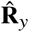 is a 180° rotation matrix. The python functions matrix_power from the numpy package and expm from the scipy package were used to calculate **M**_tot_. Chemical exchange saturation transfer (CEST) experiments were recorded using a Bruker implementation of the CEST experiment described by Vallurupalli and co-workers (2012) (21). A weak ^15^N radio frequency field of *γB*_1_ / 2*π* = 25 Hz, and a CEST exchange period *T*_ex_ of 400 ms were used. The power level for the saturation radio frequency field was calculated from the calibrated ^15^N hard pulse assuming linear amplifiers. The saturation field offset was varied from −1250 to 1250 Hz in intervals of 25 Hz covering the entire ^15^N spectral width. A reference spectrum was recorded with *T*_ex_ set to 0. CEST profiles were fitted to the Bloch-McConnell equation assuming a two-site exchange model, or to the Bloch equation, using a similar strategy as that used to compute the evolution of the magnetization during the CPMG period. The calculations considered that the intrinsic relaxation rates of CALX-CBD1 in the Apo state and in the Ca^2+^-bound state are equivalent, 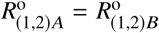. The exchange rate, *k*_ex_, the population and offset of the Apo state, *p*_*A*_ and Ω_Apo_, respectively, and the ^15^N R_1_ and R_2_ rates, were treated as free parameters. Ca^2+^ dissociation rates were obtained by doing *k*_off_ = *k*_ex_ × *p*_*A*_, where *p*_*A*_ = 1 − *p*_*B*_, where *A* refers to the population of the Apo state and *B* to the fully Ca^2+^-bound state.

The NMR titration was carried out using a sample of 180 *µ*M of CALX-CBD1 in 10 mM Tris pH 7.4, containing 200 mM NaCl, 1 mM PMSF as protease inhibitor, and 10% of D_2_O. This preparation was previously incubated with 10 mM EDTA to remove residual Ca^2+^. The EDTA was removed by a series of concentration and dilutions using an Amicon with cutoff of 3 kDa at 4 °C. A series of ^1^H− ^15^N HSQC spectra were recorded at 308 K as matrices of 1024 × 128 complex points. The NMR data was processed without apodization in order to apply the theoretical model outlined in the previous section. The data set was zero filled up to 4096 × 2048 points prior to Fourier transformation. Aliquots from a 5 M CaCl_2_ stock solution, prepared in the same buffer, were added to the NMR sample up to final Ca^2+^ concentrations of 5, 15, 25, 50, 70, 90, 190, 340, 490, 740, 990, 1490, 1990 or 3990 *µ*M. Only well isolated ^1^H− ^15^N HSQC cross peaks belonging to residues 452-456, 458-460, 479, 481, 483, 486-491, 512, 514-516, 518-520, 523, 525, 549, 550-552 and 555 of CALX-CBD1 in the Ca2+ − bound form were considered for the analysis. Cross peak intensities were determined from interpolation of a two-dimensional numerical function (2D spline) as detailed in Fig. S1 in the Supporting Material. The factor *η* was calculated according to Eq. 1 for each cross peak along the titration. Only intensities above the noise level of each spectrum were considered for the analysis. The baseline threshold was determined by calculating the absolute intensity at 1σ of the baseline distribution (68%). The intensity level at the plateau of the binding curve, *I*_max_, was determined as the mean intensity value of the three greatest Ca^2+^ concentrations: 1490, 1990 and 3990 *µ*M of CaCl_2_. The NMR titration curves were fitted to Eq. 7. The protein population in the Apo state, *p*_*A*_, was calculated according to the Hill model. The dissociation constant (*K*_*d*_) and the Hill coefficient (*n*_Hill_) were treated as free parameters. The ^1^H and ^15^N chemical shift differences between the Apo and the Ca^2+^-saturated states (ΔΩ_N_ and ΔΩ_H_), and the Ca^2+^ dissociation rates (*k*_off_), were normalized by the respective 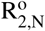 and 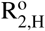 relaxation rates. We used a Gaussian distribution as prior information for the normalized dissociation rate, 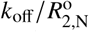. The distribution was centered at 1.34 and σ equal to 10% of this value. The prior information was based on the *k*_off_ values obtained from the CPMG relaxation dispersion experiment, and on the ^15^N transverse relaxation rates determined for CALX-CBD1 in the bound state.

### Calorimetry

Isothermal titration calorimetry (ITC) experiments were carried out using a Malvern VP-ITC instrument, at 35 °C. Samples of CALX-CBD1, of nominal concentrations 5, 10, 15 and 20 *µ*M, in 20 mM Tris-HCl pH 7.4, 200 mM NaCl, 5 mM *β*-mercaptoethanol and 1% (V/V) glycerol, were treated with 10 mM of EDTA. The EDTA was removed by buffer exchange using an Amicon centrifugation device, and the sample was transferred to the calorimeter cell. The injection syringe was filled with a solution of CaCl_2_, at 500 *µ*M, in the same buffer. The experiments consisted of a first injection of CaCl_2_ of 2 *µ*l, followed by 28 injections of 10 *µ*l of CaCl_2_. Peak integration was carried out using the instrument software, providing the heat energy released in each Ca^2+^ injection. The ITC binding curves were fitted to Eq. S28 assuming different stoichiometric binding models as described in the Supporting Material. The Bayesian approach used to fit the ITC data was based on Nguyen and co-workers (2018) (22). Fits to the Hill model considered the dissociation constant, *K*_*d*_, the Hill coefficient (*n*_Hill_), and the molar binding enthalpy (Δ*H*_*bind*_) as free parameters, while fits to the stepwise model treated *K*_*d*,1_, *K*_*d*,2_, *K*_*d*,3_, *K*_*d*,4_, and Δ*H*_*bind*_ as free parameters. As proposed by Nguyen and co-workers (2018) (22), we included a set of nuisance parameters involved in the theory, among which are the ligand concentration in the syringe [*L*]_o_, the protein concentration in the cell [*P*] _tot_, the heat offset *Q*_o_ and the Gaussian error of the ITC data, σ, for each ITC curve. The latter was considered the same for all data points. Therefore, in total there were 16 nuisance parameters for each stoichiometric model. In the case of the stepwise model, the protein concentration in the calorimeter cell was taken from a Gaussian distribution centered at calibrated values. The calibration of the protein concentration values was done by a global fitting of the ITC binding curves to a model of four independent binding sites with the same affinity. This calibration was important due to the inaccuracy on the determination of the active protein concentration. The script implemented to globally fit ITC data using Bayesian statistics is available upon request.

Micro Differential Scanning Calorimetry (*µ*DSC) experiments were performed on a VP-DSC (Microcal) instrument. The CALX-CBD1 concentration used in the DSC experiments was approximately 100 *µ*M with or without CaCl_2_ at a molar ratio of 10:1 (Ca^2+^:CBD1).

## RESULTS

### CALX-CBD1 displays significant backbone flexibility in the Apo state

The ^1^H-^15^N HSQC spectrum of the CALX-CBD1 domain recorded in the absence of Ca^2+^ displayed well-dispersed cross peaks, which are indicative of a well-folded protein. However, approximately 23 of the expected ^1^H-^15^N correlations were missing. All Apo state residues that could be assigned are located at the bottom of the domain, opposite to the four Ca^2+^-binding sites (Fig. 1). In contrast, in the presence of 20 mM of CaCl_2_ all the expected ^1^H^15^N cross peaks were observed (Fig. 1 and Fig. S2 in the Supporting Material), and 111 out of 121 backbone resonances were successfully assigned through the analysis of triple-resonance NMR experiments. These observations suggest that the tip of the domain, near the Ca^2+^-binding sites, experiences flexibility at the NMR chemical shift time scale, but becomes rigid upon Ca^2+^-binding. Consistent with this observation, the crystal structure of CALX-CBD1 in the Apo state does not show electron density for residues 516-522 at the EF loop, indicating flexibility at the Ca^2+^-binding region (15). The EF loop contains six negatively charged residues. It is, therefore, not surprising that it exhibits flexibility in the Apo state. *µ*DSC experiments showed that the binding of Ca^2+^ leads to an increase of CALX-CBD1 thermal stability by approximately 15 °C (Fig. 1). A similar increment in thermal stability upon Ca^2+^-binding was observed for classical C2 domains belonging to PKC isoenzymes (23). In addition, the width at half height, *T*_1 / 2_, decreased from 6.18 °C to 4.05 °C indicating that the ensemble of CALX-CBD1 conformations became narrower upon Ca^2+^ binding. Measurements of ^15^N relaxation rates, R_1_ and R_2_, and the heteronuclear NOE, carried out in the presence of excess of Ca^2+^, confirmed that CALX-CBD1 behaves as a rigid protein in the Ca^2+^-bound state (Fig. S3 in the Supporting Material). An analysis of the overall tumbling rate based on ^15^N R_2_/R_1_ ratios for all ^1^H^15^N spin pairs assuming isotropic tumbling, yielded *τ*_*c*_ = 9.5 ns under our NMR conditions. Larger ^15^N transverse relaxation rates were detected for residues 539-546, at *β*-strand G, near the *β*-bulge, far away from the Ca^2+^-binding sites (Fig. S3). These larger R_2_ rates could be due to a conformation exchange process at slow (*µs* to ms) timescale, or to transient oligomerization of the domain involving the *β*-bulge. Consistent with the latter hypothesis, we observed that the NMR sample tends to aggregate with time.

### CALX-CBD1 Ca^2+^-binding affinity determined by NMR

To investigate the mechanism of Ca^2+^-binding to CALX-CBD1, we carried out NMR titration experiments. ^1^H-^15^N HSQC spectra of CALX-CBD1 recorded in the absence and presence of increasing Ca^2+^ concentrations showed that Ca^2+^-binding occurs in slow exchange at the NMR chemical shift time scale. Furthermore, only cross peaks for the Apo and the Ca^2+^-saturated states were observed, indicating that four Ca^2+^ bind highly cooperatively to CALX-CBD1, as observed previously for the NCX-CBD1 (24). Ca^2+^-binding curves were calculated by monitoring the intensity increase of well isolated NMR signals of the protein in the bound state according to Eq. 1. In order to avoid bias in the determination of the cross peak intensities, a 2D cubic spline function was maximized for each cross peak as shown in Fig. S1. Interestingly, all residues located at the tip of the domain, near the Ca^2+^-binding sites, or involved in Ca^2+^ coordination such as D552, displayed sigmoidal Ca^2+^-binding curves. In contrast, cross peaks of residues located further away from the Ca^2+^-binding sites, such as A487, displayed nearly hyperbolic binding curves (Fig. 2 and Fig. S4 in the Supporting Material). As shown in (Fig. 2A), the accentuated sigmoidal Ca^2+^-binding curves did not agree with the Hill model, even when maximum cooperativity, *n*_Hill_ = 4, was assumed. Therefore, the accentuated sigmoidal profile of some of the Ca^2+^-binding curves is not due to ligand binding cooperativity. Since CALX-CBD1 experiences significant backbone flexibility in the Apo state, the binding of Ca^2+^ must be accompanied by conformational changes at the Ca^2+^-binding sites. Therefore, we examined the possibility that conformational selection or induced-fit contributions to the Ca^2+^-binding equilibrium would lead to highly sigmoidal Ca^2+^ binding curves. These possibilities were discarded because simulations of the stepwise model, considering four binding events, and a induced-fit or conformational selection step, did not lead to binding curves displaying sigmoidal behavior.

**Figure 2:**
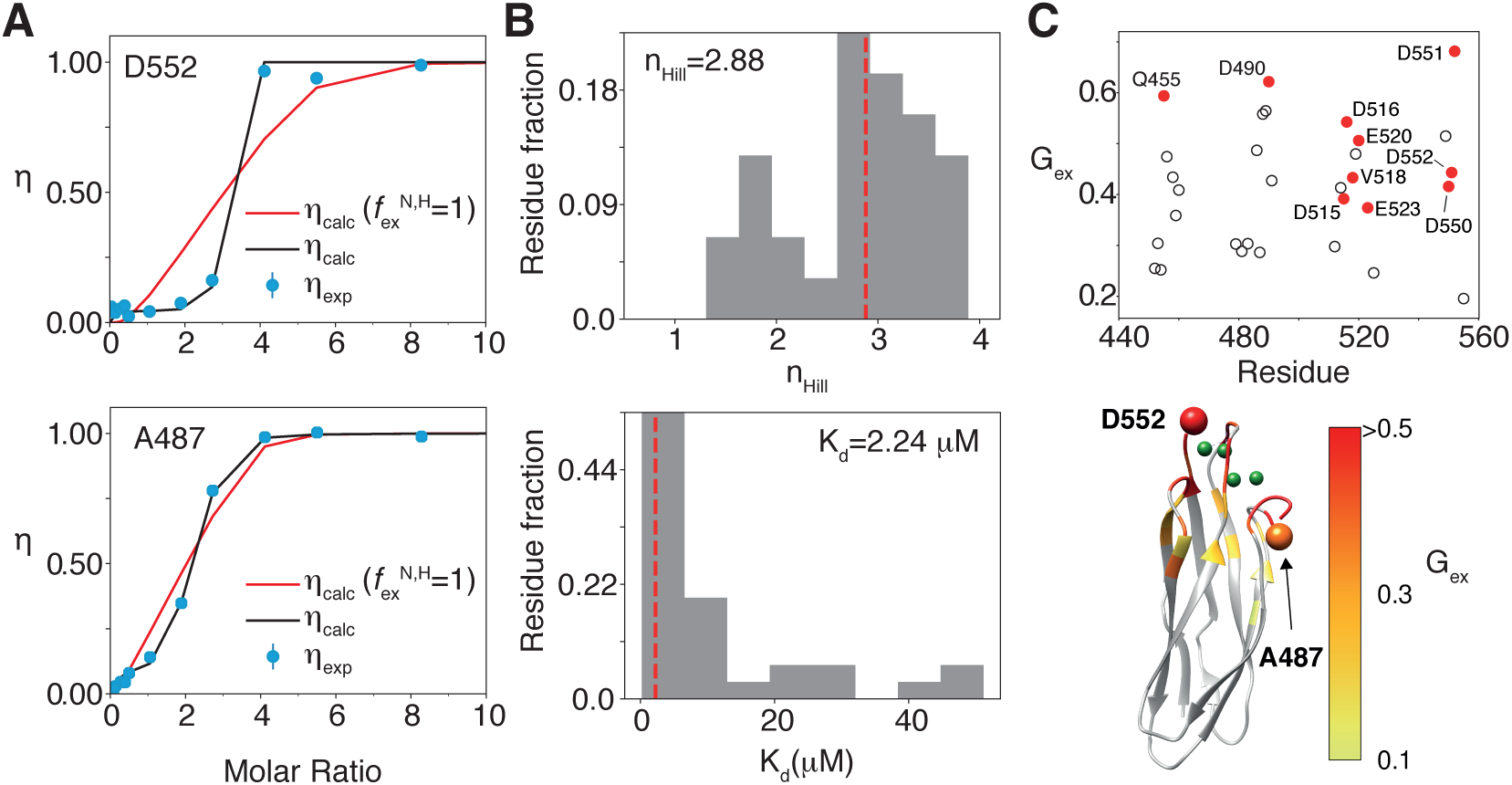
NMR Titration. A) Typical examples of NMR titration curves. Residue D552 displays an accentuated sigmoidal behavior (top left) in comparison with A487 (bottom left). Red lines are fits to the Hill model, and black lines are fits to Eq. 7 using the binding polynomial for the Hill model. B) Histograms of the obtained values for the Hill coefficient (*n*_Hill_) and affinity (*K*_*d*_) for all analyzed residues. The red line indicates the median of the distribution. C) Quantitative analysis of the magnitude of exchange contribution to the NMR titration curve as a function of the residue number, and mapped on the crystal structure of CALX-CBD1 (PDB 3EAD). The *G*_ex_ parameter is defined in Fig. S5 in the Supporting Material. Residues indicated in red (left) correspond to the ones involved in Ca^2+^ coordination.

The sigmoidal behavior of the Ca^2+^-binding curves can be explained by considering the existence of chemical exchange contributions to the NMR lineshapes. Simulations of binding curves computed with Eq. 7 and using the exact solution for 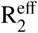, Eq. 3, in the case of a two-site chemical exchange model (16), showed that deviations from the usual behavior of the Hill model might be significant outside the extreme limits of the titration curve (Ca^2+^:CBD1≈ 0 or Ca^2+^:CBD1≈ 4), i.e. when 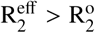 (Fig. 3). In the absence of chemical exchange, i.e. when ΔΩ = 0 or *k* _off_ = 0, *η* is equivalent to *p*_*B*_, and the Ca^2+^-binding curve follows the Hill model (Fig. 3). In contrast, when the chemical exchange contributions to R_2_ increase, the binding curve may deviate from the expected behavior as observed in our experimental data (Fig. 2). Although this simulation considered only exchange contributions to 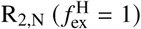, a similar effect would be observed when taking into account the simultaneous contribution of the two nuclei, ^1^H and ^15^N.

**Figure 3:**
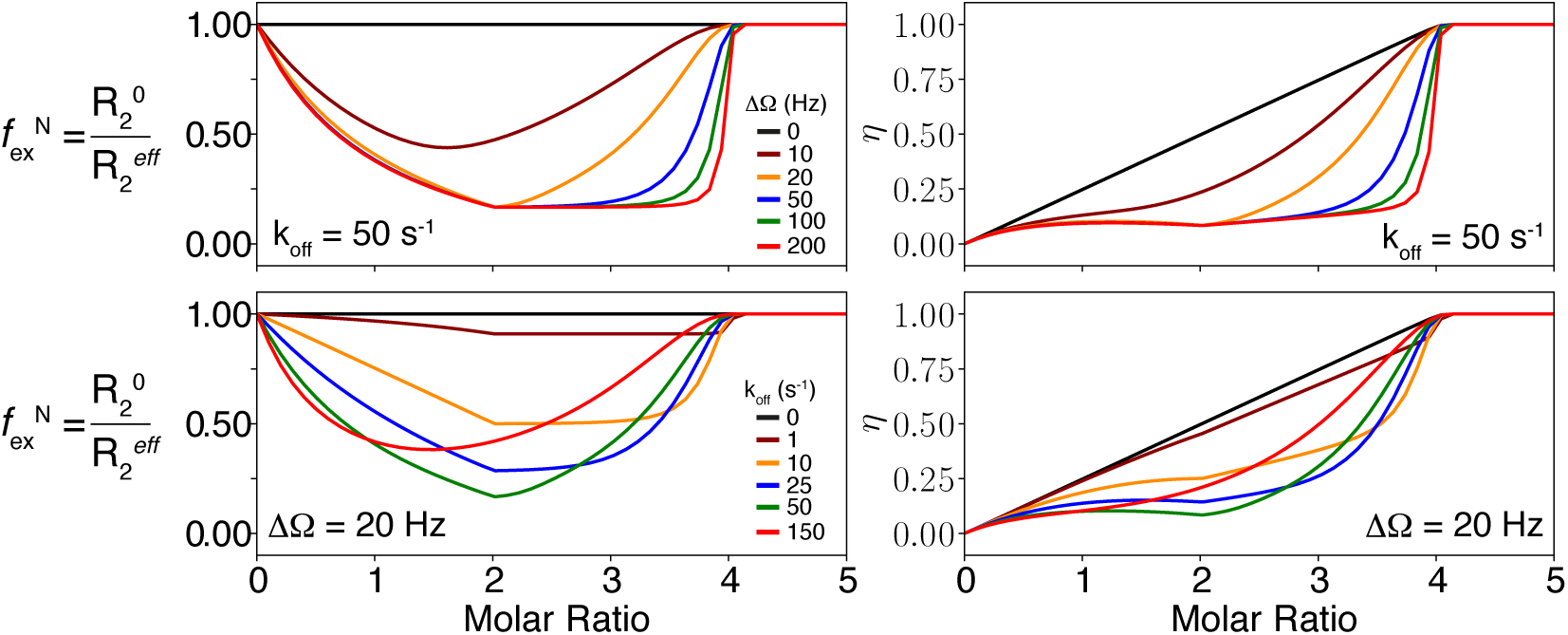
Contributions of chemical exchange to NMR-titration curves. Simulation of *η* (right) and 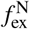 (left) as a function of the ligand:protein molar ratio. On the top panel, *k*_off_ was kept constant while ΔΩ was varied from 0 to 200 Hz. In the bottom panel, ΔΩ was left constant while *k*_off_ was varied from 0 to 150 s^−1^. The simulations assumed that ligand binding occurs according to the Hill model (Eq. S26 in the Supporting Material), and considered that 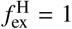. In all cases the total protein concentration was 180 *µ*M, dissociation constant *K*_*d*_ = 5*µ*M, *n*_Hill_ = 4, and 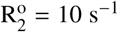.

Analysis of the probability distribution functions of the free parameters indicated that *K*_*d*_ and *n*_Hill_ are well defined for most residues. The representative values obtained from the median of the distributions of all analyzed residues are shown in Fig. 2B, and indicate that *n*_Hill_ = 2.88 and *K*_*d*_ = 2.24 *µ*M. The determined Hill coefficient (*n*_Hill_ = 2.88) indicates that Ca^2+^ binds cooperatively to CALX-CBD1. Importantly, this analysis did not include any prior for the chemical exchange variables 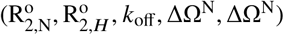. The extent of deviation from the profile expected for each binding curve in the absence of exchange was estimated through an empirical parameter *G*_ex_. The latter was defined as the ratio of the area of the 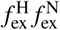 function relative to the area of a rectangle from molar ratios 0 to 4 (Fig. S5 in the Supporting Material). This analysis does not depend on the convergence of the fitted parameters, but rather depends on the shape of the binding curve. *G*_ex_ may be qualitatively associated to the degree of chemical exchange contribution to the transverse relaxation rates at a specific HN site. As shown in Fig. 2C, all residues involved in Ca^2+^ coordination in the crystal structure, displayed the largest deviations, hence, the largest *G*_ex_ values.

### Determination of Ca^2+^ dissociation rates (*k*_off_) by ^15^N relaxation experiments

Chemical exchange saturation transfer (CEST) experiments are informative about dynamic processes that occur in the slow exchange regime, at the millisecond time scale (21). Therefore, we investigated whether ^15^N CEST experiments could reveal information on the kinetics of Ca^2+^-binding to CALX-CBD1. The observed CEST effects were interpreted assuming a two-site exchange model, corresponding to CALX-CBD1 in equilibrium between the Apo and the fully Ca^2+^-saturated states. In order to take advantage of all backbone resonance assignments available for CALX-CBD1 in the Ca^2+^-bound state, we recorded ^15^N-CEST experiments on samples prepared in the presence of CaCl_2_ at the stoichiometric 4:1 molar ratio (Ca^2+^:CALX-CBD1). Under this condition, cross peaks due to the Apo state are not detected. As shown in Fig. 4, three different CEST profiles were observed. Residues located at the bottom of the domain, i.e. far away from the Ca^2+^-binding sites, displayed a single and narrow CEST dip, which did not change upon the addition of excess of Ca^2+^ (Fig. 4). In contrast, residues at the tip of the domain, or involved in Ca^2+^ coordination, displayed broadened CEST dips, which became significantly narrower upon the addition of excess of Ca^2+^. A total of four residues located at or near the Ca^2+^-binding sites, N456, V518, F519, and L549, displayed a single CEST dip and a shoulder, while only G550, which coordinates Ca^2+^ at Ca3, displayed two well-separated CEST dips. Residues that displayed the greatest broadening effects are the ones closer to the Ca^2+^-binding sites, and are expected to experience the largest chemical shift changes upon Ca^2+^-binding (Fig. 4). The absence of two well separated CEST dips precluded a quantitative analysis of the ^15^N-CEST profiles for most CALX-CBD1 residues, with the exception of N456, V518, F519, and G550. Their ^15^N-CEST profiles were fitted to the Bloch-McConnell equation assuming a two-site exchange model. It was found that the population of the Apo state, *p*_*A*_, was in the range of 6 - 8 %, while *k*_ex_ in the range of 240 - 480 *s*^−1^ (Fig. S6 in the Supporting Material).

**Figure 4:**
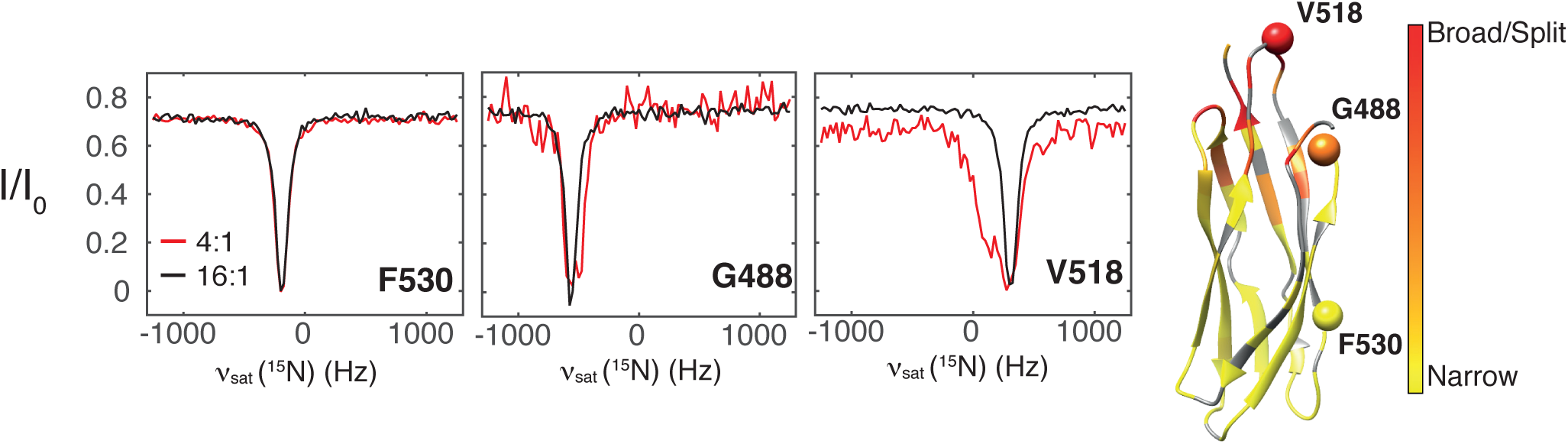
Representative ^15^N CEST profiles. Peaks that show no evidence of chemical exchange (Narrow) at the molar ratio of 4:1 (Ca^2+^:CALX-CBD1), that show nearly two CEST dips (Broad) or eventually two dips (Split), are color coded on the crystal structure of CALX-CBD1 (PDB 3EAD (15)). The color coding was based on the ratios of ^15^N R_2_/R_1_ obtained at 4:1 molar ratio (Ca^2+^:CBD1) relative to 16:1. The relaxation rates were obtained by fitting the CEST profiles to the Bloch equation. Residues 530, 488 and 518 are shown as spheres.

To obtain further kinetic information on the binding of Ca^2+^ to CALX-CBD1, we recorded CPMG based ^15^N R_2_ dispersion experiments on CALX-CBD1 samples prepared in the presence of CaCl_2_ at the stoichiometric 4:1 molar ratio (Fig. 5). As observed in the CEST experiment, residues located near the Ca^2+^-binding sites displayed more dispersion than the ones located further away. The dispersion was significantly reduced or absent in the presence of an excess of Ca^2+^, indicating that the exchange phenomenon was due to the Ca^2+^-binding equilibrium (Fig. 5). Dispersion curves obtained at two spectrometer frequencies (600 and 800 MHz) were fitted simultaneously to a two-site exchange model, yielding the absolute chemical shift difference between the Apo and the Ca^2+^-bound states, the exchange rate, and the population of the Ca^2+^-bound state. Prior information on *k*_ex_ and *p*_*A*_ obtained from the quantitative analysis of the CEST experiment were used to fit the CPMG dispersion curves. Hence, sampled *k*_ex_ and *p*_*B*_ values were taken from Gaussian distributions centered at 350 s^−1^ with 1σ = 150 s^−1^, or centered at 95% with 1σ = 5%, respectively. The dispersion curves of a total of seventeen residues could be analysed, yielding a representative Ca^2+^ dissociation rate (*k*_off_) of ∼ 19.96 s^−1^, and the Apo state population of ∼ 4.5% (Fig. 5). These numbers are in good agreement with the Ca^2+^ dissociation rate of 18.5 s^−1^ measured by stopped flow using a fluorescent Ca^2+^ chelator (13).

**Figure 5:**
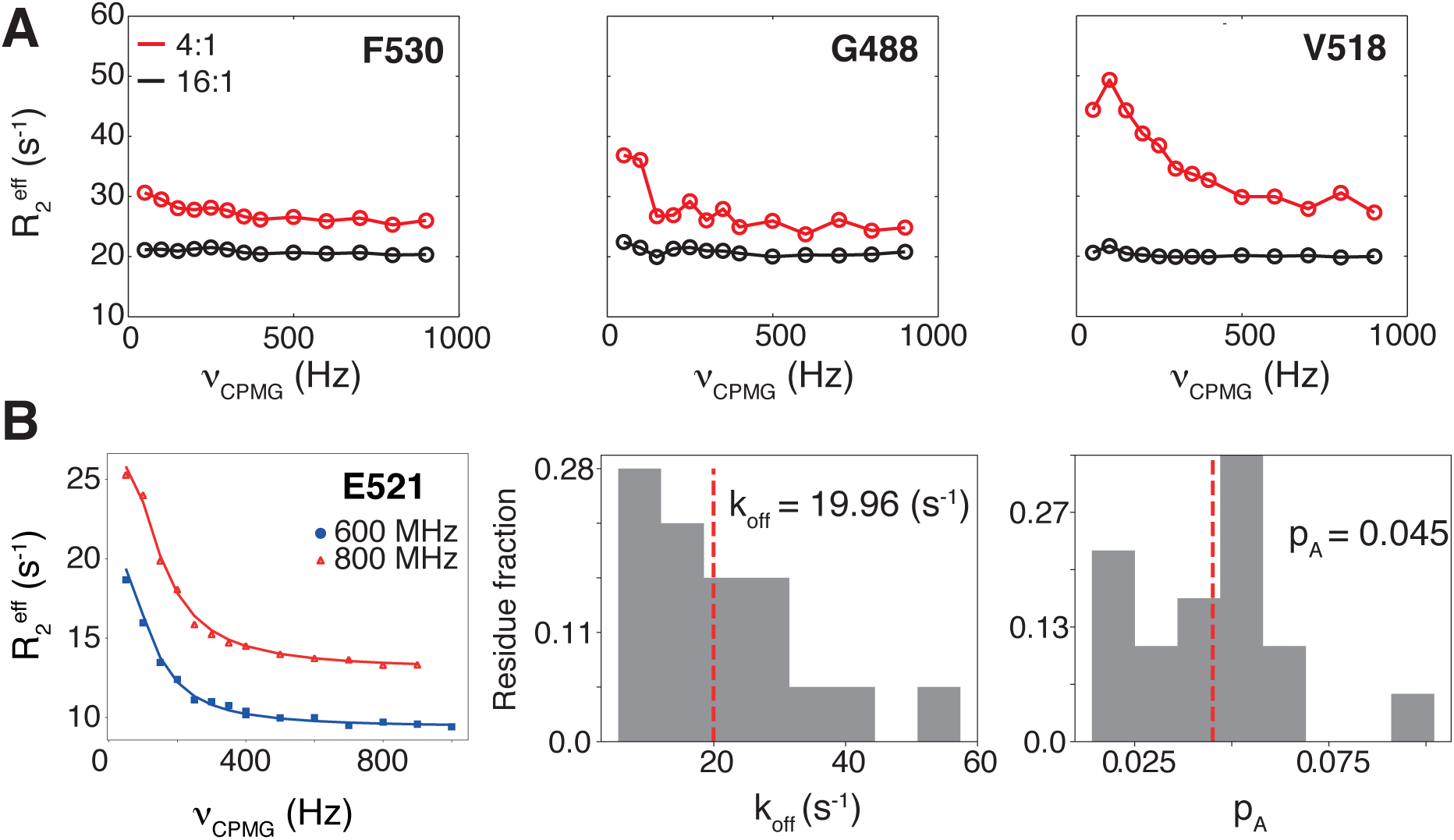
^15^N relaxation dispersion experiments. A) Representative ^15^N CPMG relaxation dispersion curves obtained at 4:1 (red) and 16:1 (black) Ca^2+^:CALX-CBD1 molar ratios. The residue number is indicated in the figure, and the location on the structure is shown in Fig. 4. A constant relaxation time of 50 ms was used for these experiments. B) Fitting of ^15^N CPMG relaxation dispersion curves obtained for E521, obtained at 800 MHz and 600 MHz (^1^H frequency) using a constant relaxation time of 40 ms, to a two-site exchange model (left). Histograms of the Ca^2+^-dissociation rates and populations of the Apo state determined by fitting the profiles obtained for residues 452-453, 455-456, 481, 488, 490-491, 515-516, 518-519, 521, 523, 549, 552, 555. The red line indicates the median of the distribution, while the representative values are indicated in the figure.

### Thermodynamics of Ca^2+^-binding

The thermodynamics of Ca^2+^-binding to CALX-CBD1 was investigated by ITC. In order to have a better statistics for the fitted parameters, four ITC experiments were carried out using different protein concentrations in the calorimeter cell. Global fitting of all ITC binding curves to a stoichiometric model assuming four independent binding sites with the same affinity (*n* = 4), yielded a dissociation constant, *K*_*d*_ = 0.214 ± 0.005 *µ*M, in agreement with previous ITC data reported in the literature (14). However, since CALX-CBD1 binds Ca^2+^ cooperatively, we tested other stoichiometric models. Hence, we fitted the ITC binding curves to the Hill model (Fig. S7 in Supporting Material), obtaining a dissociation constant of *K*_*d*_ = 2.59*µ*M, and a Hill coefficient, *n*_Hill_ = 3.07 (Table 1), in excellent agreement with the *K*_*d*_ and *n*_Hill_ determined independently by NMR titrations (Fig. 2). Since a characteristic of the Hill model is the strong correlation between the Hill coefficient (*n*_Hill_) and the binding affinity (*K*_*d*_) (Fig. S7), we tested whether a stepwise model of four binding events was able to capture the Ca^2+^-binding cooperativity of CALX-CBD1. We assumed a mean molar binding enthalpy (Δ*H*_bind_, see Eq. S31 in the Supporting Material) instead of individual binding enthalpies for each Ca^2+^-binding site. This assumption is justified because the four Ca^2+^ binding sites are located at approximately 4Å from each other in the CALX-CBD1 crystallographic structure (15), and each Ca^2+^ ion is coordinated by two or more electron donor groups from different amino acids. Here, the simultaneous fitting of the four ITC binding curves was key to alleviate the degeneracy among the stepwise dissociation constants (Fig. 6). In addition, the application of Baysean statistics to sample the space of the parameters was critical to make reliable estimations for *K*_*d*,1_ − *K*_*d*,4_, and to examine the degree of correlation between each of them (Fig. 6). The probability distribution functions indicated that *K*_*d*,3_ and *K*_*d*,4_ are well defined and equal to 0.34 and 0.85 *µ*M, respectively (Fig. 6, Table 1). *K*_*d*,1_ is well defined with a median value of 148 *µ*M but its distribution is rather wide. Finally, the degeneracy of *K*_*d*,2_ was only partially resolved. Nevertheless, the trend for *K*_*d*,2_ is in the nM range (Fig. 6, Table 1). The greater degeneracy observed for *K*_*d*,2_ relative to the other dissociation constants could be related to the fact that the experiments were carried out using protein concentrations well above its value.

**Table 1:** Thermodynamic binding constants obtained using ITC. *K*_*d*_ as well as *K*_*d,i*_ are in units of *µ*M and Δ*H*_*bind*_ in kcal/mol. *β*_*i*_ is in units of *µ*M^−*i*^. *ρ* is a dimensionless factor.

| | $K_d$ | $n$ | $K_{d,1} (\beta_1)$ | $K_{d,2} (\beta_2)$ | $K_{d,3} (\beta_3)$ | $K_{d,4} (\beta_4)$ | $\rho_2$ | $\rho_3$ | $\rho_4$ | $\Delta H_{bind}$ |
| --- | --- | --- | --- | --- | --- | --- | --- | --- | --- | --- |
| Hill | 2.59 | 3.07 | — | — | — | — | — | — | — | -5.42 |
| Stepwise | — | — | 148 (0.0068) | 0.00023 (29.4) | 0.34 (86.4) | 0.85 (101.7) | 1310 | 1.26 | 1.14 | -4.99 |

**Figure 6:**
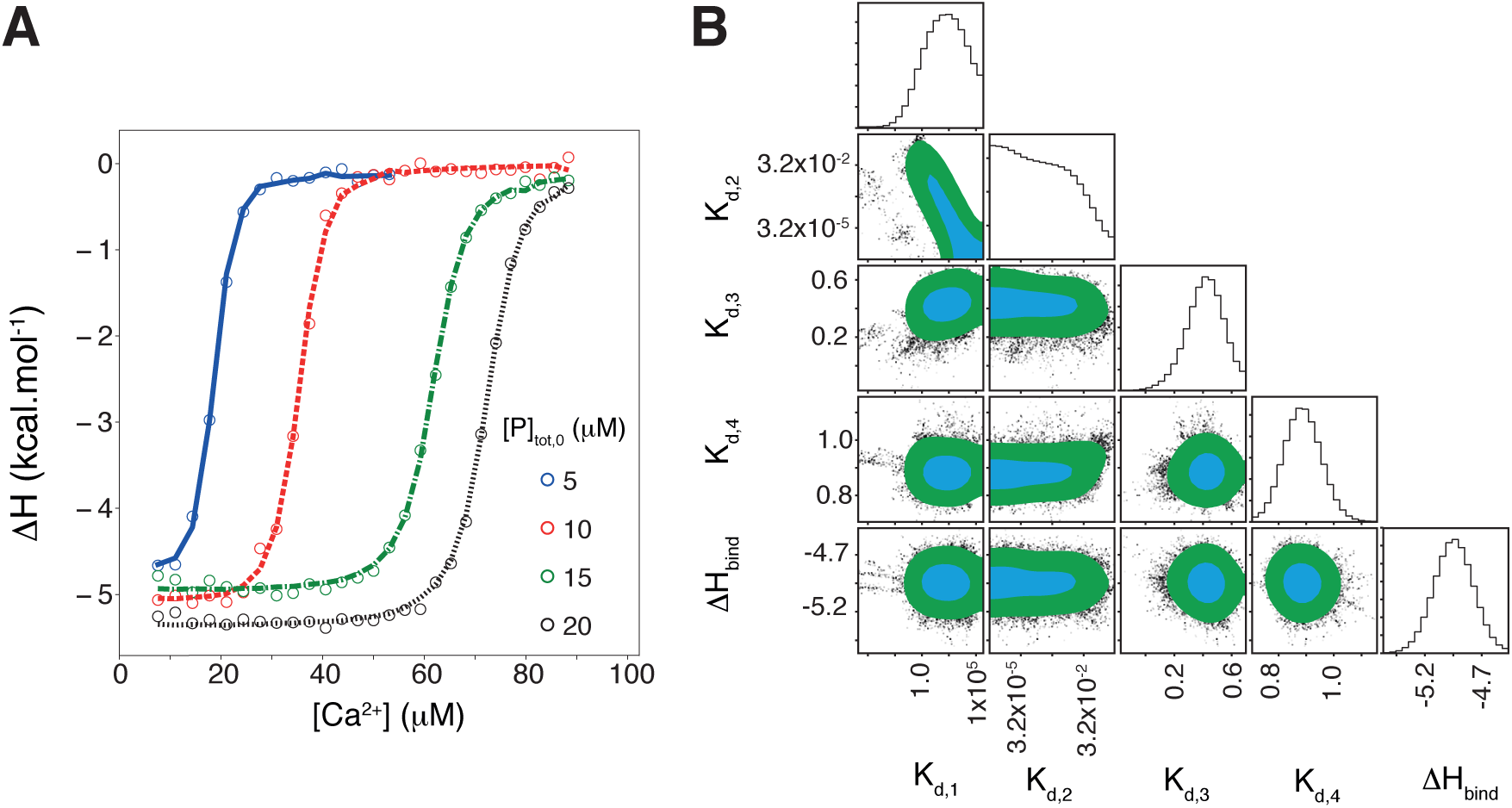
Ca^2+^-binding cooperativity investigated by ITC. A) Calorimetric titrations of CALX-CBD1 with CaCl_2_ using different total protein concentrations in the calorimeter cell ([*P*] _tot,0_). The thermograms were fitted to the stepwise model assuming four binding events. B) One dimensional marginalized parameter distributions and 2D marginalized distributions of pairs of thermodynamic parameters. The 1s confidence contour region is shown in cyan, while the 2s confidence contour region is shown in green plus cyan. *K*_*d,i*_ are in units of *µ*M and Δ*H*_*bind*_ in kcal/mol. The plots are in logarithmic scale.

In order to evaluate the Ca^2+^-binding cooperativity of CALX-CBD1 according to the stepwise model, we calculated the *ρ* factor as proposed by Wyman and Phillipson (1974) (25). The latter is defined as the ratio between the overall association constants *β*_*i*_ (see Eq. S19 in the Supporting Material) normalized by the number of combinations in each binding event (*W*_*n,i*_) (25, 26):

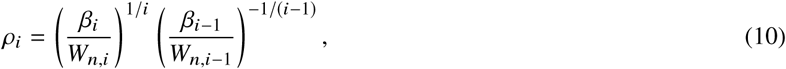

where *i* = 2, …, *n*, and *n* is the total number of binding sites. When *ρ*_*i*_ > 1 the binding sites display positive cooperativity. In contrast, when *ρ*_*i*_,,; 1 the binding sites are anticooperative or independent (25). As shown in Table 1, binding of two, three or four Ca^2+^ ions to CALX-CBD1 is favored by at least three orders of magnitude in comparison with the binding of a single Ca^2+^ ion, indicating a large positive cooperativity among the four Ca^2+^-binding sites.

## DISCUSSION

In this study we investigated the Ca^2+^-binding properties of CALX-CBD1 using NMR spectroscopy and isothermal titration calorimetry experiments, in combination with Bayesian statistics for data analysis. The NMR data indicated that CALX-CBD1 binds four Ca^2+^ in an all-or-none fashion, as observed previously for the NCX-CBD1 (24, 27). Furthermore, indirect measurements of Ca^2+^ dissociation rates, obtained from the analysis of ^15^N R_2_ dispersion and ^15^N CEST experiments, are in agreement with the Ca^2+^ off-rates determined directly by stopped-flow fluorescence measurements (13). The NMR spectra showed no evidence of the existence of an intermediate state of CALX-CBD1 with Ca^2+^ at sites Ca1 and Ca2, as observed by crystallography (15). A possible explanation for this discrepancy is that, in highly cooperative systems, the intermediate states are less populated than the Apo and the fully bound states.

It is noteworthy that the binding affinity (*K*_*d*_) and the *n*_Hill_ coefficient determined by the analysis of NMR and ITC binding curves were in very good agreement with each other (Figure 2 and Table 1). The determined *n*_Hill_ indicates that the binding of four Ca^2+^ to CALX-CBD1 is a highly cooperative process (Figure 2). Curiously, the *n*_Hill_ coefficient obtained for CALX-CBD1 in this study (*n*_Hill_ ≈ 3.0) is greater than that determined for the NCX-CBD1 from the analysis of equilibrium ^45^Ca^2+^ binding assays (*n*_Hill_ = 2.0) (28), suggesting that the CALX-CBD1 binds Ca^2+^ slightly more cooperatively than its mammalian orthologue. Since the two CBDs share the same Ca^2+^ binding mode, the reason for this difference is not clear.

Even though the Hill model captures the binding cooperativity of the system in terms of the Hill coefficient, it represents an extreme limiting situation where all ligands bind simultaneously. Moreover, the Hill model has the inconvenience that the *n*_Hill_ is strongly correlated with the *K*_*d*_ as indicated by our analysis (Fig. S7). Therefore, in order to have additional insights on the cooperative binding of Ca^2+^ to CALX-CBD1, we analysed the ITC binding curves assuming a stepwise model of four binding events. Ca^2+^-binding cooperativity of CALX-CBD1 was evaluated in terms of the *ρ* factor. According to Wyman and Phillipson (25), the *ρ*_*i*_ factor may be interpreted as the ratio of the arithmetic mean probability of binding *i* and *i* + 1 ligands, without distinguishing the binding sites. Then, cooperativity is understood as the excess probability of binding *i* + 1 ligands over the probability of binding *i* ligands. This analysis showed that the probability of binding of two, three or four Ca^2+^ ions to CALX-CBD1 is at least three orders of magnitude greater than the probability of binding of a single Ca^2+^ (*ρ*_2_ = 1310, while *ρ*_3,4_ = 1.1 − 1.3) (Table 1).

Interestingly, the NMR titration curves of residues located at short distances from the Ca^2+^-binding sites, or involved in Ca^2+^ coordination, displayed significant deviations from the expected behavior of a typical binding curve following the Hill model (Fig. 2C). Such deviations were explained by considering chemical exchange contributions to the cross peak intensities. This interpretation is consistent with the observation that residues whose binding curves are more sigmoidal map to the same region of the structure as the ones displaying the largest ^15^N chemical shift differences (ΔΩ_N_) between the Ca^2+^-bound and Apo states, as determined from the analysis of ^15^N R_2_ dispersion curves (Fig. S8 in the Supporting Material and Fig. 2C). It is curious, however, that the NMR titration curves obtained for the NCX-CBD1 domain in the NCX-CBD12 construct were all nearly hyperbolic (27). The explanation for this discrepancy is not clear, but the deviations from the expected behavior depend in a complex way on the exchange rates and on the chemical shift differences as shown in Fig. 3.

## CONCLUSIONS

Na^+^/Ca^2+^ exchangers play critical roles for the maintenance of the intracellular Ca^2+^ homeostasis. The basal intracellular Ca^2+^ concentration in *Drosophila* photoreceptor cells is approximately 0.16 *µ*M, but it may be elevated to 200 *µ*M very rapidly after light stimulation (29). In this context, the exchanger activity must be tightly regulated in order to meet the cell’s needs. Activation of the NCX by intracellular Ca^2+^ is a highly cooperative process (30). The efficient Ca^2+^-regulation of the exchangers must be, at least in part, explained by the cooperative binding of Ca^2+^ to their CBD domains. Therefore, in this study we revisited the cooperativity of Ca^2+^-binding to CALX-CBD1, the only domain that has binding sites for regulatory Ca^2+^ in the *Drosophila* Na^+^/Ca^2+^ exchanger. We found that four Ca^2+^ ions bind in a highly cooperative fashion to CALX-CBD1. According to the stepwise model, the binding affinity of the first Ca^2+^ is of the order of 150 *µ*M, while the binding affinities of the second, third and fourth Ca^2+^ are in the nM and *µ*M ranges, respectively. This observation indicates that the probability of binding of a single Ca^2+^ is significantly lower than that of binding from two up to four Ca^2+^ ions. Ca^2+^ dissociation must also be highly cooperative as indicated by NMR, with off-rates of the order of 20 s^−1^. These findings provide new insights on the Ca^2+^-binding cooperativity of CALX-CBD1, which is key to enable this exchanger to efficiently respond to changes in the intracellular Ca^2+^-concentration in sensory neuronal cells.

## AUTHOR CONTRIBUTIONS

All authors contributed to the study conception and design. Phelipe M. Vitale and Layara A. Abiko prepared protein samples. Layara A. Abiko assigned backbone resonances. Phelipe M. Vitale, Marcus V. C. Cardoso and Maximilia Degenhardt did ITC experiments. Preliminary analysis codes for ITC and NMR titration curves were performed by Maximilia Degenhardt. The final codes for data analysis on NMR and ITC experiments were designed and developed by Jose D. Rivera and Roberto K. Salinas. Marcus V. C. Cardoso and Layara A. Abiko acquired and processed NMR relaxation and titration data. The first draft of the manuscript was written by Roberto K. Salinas, Marcus V. C. Cardoso, Phelipe M. Vitale and Jose D. Rivera. All authors commented on previous versions of the manuscript. All authors read and approved the final manuscript.

## ACKNOWLEDGMENTS

This work was supported by grants from the São Paulo Research Foundation (FAPESP 2016/07490-1) and Conselho Nacional de Desenvolvimento Científico e Tecnológico (CNPq 420490/2016-7). MVCC (2016/17375-5), JDRE (2018/21450-8) and PMV (2017/05614-8) received FAPESP Post-Doctorate and PhD fellowships. LAA and MFSD received PhD fellowships from Coordenação de Aperfeiçoamento de Pessoal de Nível Superior (CAPES, #33002010017P0 and #88882.160170/2014-01). RKS receives a CNPq research fellowship (#311807/2017-8). The *CALX1.1* cDNA was obtained from the Drosophila Genomics Resource Center, supported by NIH grant 2P40OD010949. The authors acknowledge Bruker BioSpin for providing the pulse sequence to record ^15^N CEST experiments, the Multiuser Center for Biomolecular Innovation of the UNESP (EMU-FAPESP 2009/53989-4) for providing access to the 600 MHz NMR instrument, the Analytical Instrumentation Center of the University of São Paulo for providing access to the 800 MHz NMR instrument, and FAPESP for providing financial resources for the acquisition of the ITC instrument (2017/06394-1).

## SUPPORTING CITATIONS

References (31–36) appear in the Supporting Material.

