## Supporting Material for "CALX-CBD1 Ca^2+^-binding cooperativity studied by NMR spectroscopy and ITC with Bayesian statistics"

### BINDING POLYNOMIAL FORMALISM

The binding polynomial formalism (1–3) was used in order to derive expressions for the stoichiometric models used to analyze ITC and NMR titration data. Briefly, consider a system given by a protein $P$ with $n$ liganded states for a ligand $L$ (i.e. the protein molecule has $n$ binding sites), where the ensemble of states is in equilibrium. The concentration of protein molecules in the $i$-liganded state is defined as $[PL_i]$, then the total concentration of protein is:

$$[P]_{\text{tot}} = \sum_{i=0}^{n} [PL_i], \tag{1}$$

where $[PL_0] = [P]$ is the protein concentration in the Apo state. The mean concentration of bound ligands is $i[PL_i]$, therefore the total ligand concentration is given by:

$$[L]_{\text{tot}} = [L] + \sum_{i=0}^{n} i[PL_i], \tag{2}$$

where $[L]$ is the free-ligand concentration. The probability weight for a $i$-liganded state (i.e. the probability that a protein molecule is at the $i$-liganded state), $f_{PL_i}$, is equal to the fraction of protein molecules in that state, thus

$$f_{PL_i} = \frac{[PL_i]}{[P]_{\text{tot}}} = \frac{[PL_i]}{\sum_{i=0}^{n}[PL_i]}. \tag{3}$$

where $n$ is the maximum number of binding sites in the protein $P$. Since the states are in thermodynamic equilibrium, the concentration for a complex $PL_i$ is given by:

$$[PL_i] = \beta_i [P][L]^i, \tag{4}$$

where $\beta_i$ is the overall association constant, by definition $\beta_0 = 1$. By using the above equation, the expression (3) may be rewritten as:

$$f_{PL_i} = \frac{\beta_i [L]^i}{\sum_{i=0}^{n} \beta_i [L]^i} = \frac{1}{Z}\beta_i [L]^i. \tag{5}$$

Here the function $Z$ is the so called binding polynomial and it is defined as:

$$Z \equiv \sum_{i=0}^{n} \beta_i [L]^i = \frac{1}{[P]} \sum_{i=0}^{n} [PL_i]. \tag{6}$$

The average number of bound ligands per protein molecule, $\nu$, during a titration can be computed using the probability weight defined in Eq. 3, then

$$\langle i \rangle = \nu = \sum_{i=1}^{n} i f_{PL_i} = \frac{[L]}{Z} \frac{\mathrm{d}Z}{\mathrm{d}[L]}. \tag{7}$$

Usually the experimental data provides us the average number of ligands bound and the total ligand concentration $[L]_{\text{tot}}$. Then, the relation between free and total ligand concentrations is given by:

$$([L]_{\text{tot}} - [L]) = \sum_{i=1}^{n} i\,[PL_i] = [P]_{\text{tot}} \frac{\mathrm{d}\ln Z}{\mathrm{d}\ln[L]}. \tag{8}$$

In order to determine $\nu$ and the free ligand concentration, it is necessary to know the binding polynomial, which depends on the considered model.

#### $n$-Independent binding sites model

Consider a protein that has $n$ independent ligand binding sites, that is, ligand-binding to one site has not effect on the binding affinity of the other sites. In this case, the binding polynomial, Eq. 6, is the product of the binding polynomial of individual binding events (1, 2)

$$Z = \prod_{i=1}^{n}(1 + K_i[\mathrm{L}]), \tag{9}$$

where $i$ refers to a specific binding site with affinity $K_i$. By using Eq. 7, $\nu$ is given by:

$$\nu = \sum_{i=1}^{n} \frac{K_i[\mathrm{L}]}{1 + K_i[\mathrm{L}]}. \tag{10}$$

Here, the free ligand is related to the total ligand via:

$$([L]_{\text{tot}} - [L]) = [P]_{\text{tot}} \sum_{i=1}^{n} \frac{[L]K_i}{1 + K_i[L]}. \tag{11}$$

For $n = 4$, such as the CALX-CBD1 and $Ca^{2+}$ system, Eq. 10 allows us to compute the average number of bound ligands per protein molecule, which is given by:

$$\nu = [\mathrm{Ca}^{2+}] \left( \frac{K_1}{1 + [\mathrm{Ca}^{2+}]K_1} + \frac{K_2}{1 + [\mathrm{Ca}^{2+}]K_2} + \frac{K_3}{1 + [\mathrm{Ca}^{2+}]K_3} + \frac{K_4}{1 + [\mathrm{Ca}^{2+}]K_4} \right). \tag{12}$$

By using Eq. 11 and some basic algebra, we obtain the free ligand concentration,

$$\begin{aligned} &K_d[\mathrm{Ca}^{2+}]^5 + \left(K_d(4[P]_{\text{tot}} - [\mathrm{Ca}^{2+}]_{\text{tot}}) + K_c\right)[\mathrm{Ca}^{2+}]^4 + \left(K_c(3[P]_{\text{tot}} - [\mathrm{Ca}^{2+}]_{\text{tot}}) + K_b\right) \\ &[\mathrm{Ca}^{2+}]^3 + \left(K_b(2[P]_{\text{tot}} - [\mathrm{Ca}^{2+}]_{\text{tot}}) + K_a\right)[\mathrm{Ca}^{2+}]^2 + \left(K_a([P]_{\text{tot}} - [\mathrm{Ca}^{2+}]_{\text{tot}}) + 1\right)[\mathrm{Ca}^{2+}] \\ &- [\mathrm{Ca}^{2+}]_{\text{tot}} = 0, \end{aligned}$$

where $K_a = K_1 + K_2 + K_3 + K_4$, $K_b = K_1K_2 + K_1K_3 + K_1K_4 + K_2K_3 + K_2K_4 + K_3K_4$, $K_c = K_1K_2K_3 + K_1K_2K_4 + K_1K_3K_4 + K_2K_3K_4$ and $K_d = K_1K_2K_3K_4$.

If we assume that the $n$ independent sites have equal binding affinity, $K$, and using Eq. 9, the binding polynomial is given by:

$$Z = (1 + K[L])^n = \sum_{r=0}^{n} W_{n,r} K^r [L]^r, \tag{13}$$

where $W_{n,r}$ is the number of combinations,

$$W_{n,r} = \frac{n!}{r!(n-r)!}. \tag{14}$$

Note that for each $i$-liganded protein species $PL_i$ the observed macroscopic binding constants will depend on the number of combinations necessary to accommodate $r$ ligands into $n$ binding sites. According to the Eq. 10 the fraction of ligands per protein molecule is

$$\nu = \frac{nK[L]}{1 + K[L]}, \tag{15}$$

and the free ligand concentration is obtained via Eq. 11,

$$K[L]^2 + ((n[P]_{\text{tot}} - [L]_{\text{tot}})K + 1)\,[L] - [L]_{\text{tot}} = 0. \tag{16}$$

The above expression is a second order polynomial, where its solution is given by:

$$[L] = \frac{([L]_{\text{tot}} - n[P]_{\text{tot}})\,K - 1 + \sqrt{[(n[P]_{\text{tot}} - [L]_{\text{tot}})\,K + 1]^2 + 4K[L]_{\text{tot}}}}{2K}. \tag{17}$$

Here only the positive solution was considered.

#### Stepwise binding events

In the stepwise binding model, $n$ ligand molecules bind to the protein $P$ in a sequential order:

$$P + L \rightleftharpoons PL$$
$$PL + L \rightleftharpoons PL_2$$
$$\vdots$$
$$PL_{n-1} + L \rightleftharpoons PL_n$$

The binding polynomial for this "sequential" binding equilibrium is given by:

$$Z = 1 + \sum_{i=1}^{n} [L]^i \beta_i, \tag{18}$$

where

$$\beta_i = \prod_{j=1}^{i} K_j, \quad \beta_0 = 1. \tag{19}$$

Here, $K_i$ is the association constant of the $i$-ligand binding event. Hence, the average number of ligands bound, $\nu$, is:

$$\nu = \frac{1}{Z}\sum_{i=1}^{n}\left(i[L]^i\beta_i\right). \tag{20}$$

According to this definition, $\nu$ will vary from 0 when no ligand is bound, to $n$ when all binding sites are occupied. A general polynomial expression for the free ligand concentration, $L$, was derived from Eq. 8 and assuming the current form of $Z$, Eq. 18:

$$[L]^{n+1}\beta_n + \sum_{i=1}^{n}[L]^i\left(\beta_i\left(i[P]_{\text{tot}} - [L]_{\text{tot}}\right) + \beta_{i-1}\right) - [L]_{\text{tot}} = 0, \tag{21}$$

which allows one to calculate the free ligand concentration, $[L]$, at any point of the titration and, hence, $\nu$.

The expressions derived above are generic. In our case of interest that involves four ligand binding sites, $n = 4$, following Eq. 18 we find:

$$Z = 1 + K_a[Ca^{2+}] + K_b[Ca^{2+}]^2 + K_c[Ca^{2+}]^3 + K_d[Ca^{2+}]^4, \tag{22}$$

where $K_a = K_1$, $K_b = K_1K_2$, $K_c = K_1K_2K_3$, $K_d = K_1K_2K_3K_4$ are the overall association constants and $K_1$, $K_2$, $K_3$, and $K_4$ are the stepwise association constants of four ligands $L$ to the protein $P$. The average number of bound ligands per protein molecule, $\nu$, may be calculated applying Eq. 20:

$$\nu = \frac{K_a[Ca^{2+}] + 2K_b[Ca^{2+}]^2 + 3K_c[Ca^{2+}]^3 + 4K_d[Ca^{2+}]^4}{1 + K_a[Ca^{2+}] + K_b[Ca^{2+}]^2 + K_c\left[Ca^{2+}\right]^3 + K_d[Ca^{2+}]^4}, \tag{23}$$

and the free $[Ca^{2+}]$ is obtained applying Eq. 21:

$$\begin{aligned} &K_d[Ca^{2+}]^5 + \left(K_d(4[P]_{\text{tot}} - [Ca^{2+}]_{\text{tot}}) + K_c\right)[Ca^{2+}]^4 + \left(K_c(3[P]_{\text{tot}} - [Ca^{2+}]_{\text{tot}}) + K_b\right) \\ &[Ca^{2+}]^3 + \left(K_b(2[P]_{\text{tot}} - [Ca^{2+}]_{\text{tot}}) + K_a\right)[Ca^{2+}]^2 + \left(K_a([P]_{\text{tot}} - [Ca^{2+}]_{\text{tot}}) + 1\right)[Ca^{2+}] \\ &- [Ca^{2+}]_{\text{tot}} = 0. \end{aligned} \tag{24}$$

One should note that the above equation is equivalent to the equation for the free ligand concentration in the independent case, however $K_a$, $K_b$, $K_c$ and $K_d$ are different.

#### The Hill model

Consider a particular case of a protein $P$ that binds $n$ ligand molecules in all-or-none fashion:

$$P + n\,L \leftrightharpoons PL_n.$$

This is the Hill model. The binding polynomial, $Z$, for the Hill model is given by:

$$Z = 1 + K[L]^n, \tag{25}$$

and, hence, the average number of occupied sites, $\nu$, is obtained from Eq. 7 (1):

$$\nu = \frac{nK[L]^n}{1 + K[L]^n}, \tag{26}$$

in which $K$ is the association constant to the power of $n$ ($K = K_a^n$), $n$ is the Hill coefficient ($n_{\mathrm{Hill}}$), and $[L]$ is the free ligand concentration. The free ligand concentration as a function of the total ligand concentration is obtained applying Eq. 8:

$$K[L]^{n+1} + K(n[P]_{\mathrm{tot}} - [L]_{\mathrm{tot}})[L]^n + [L] - [L]_{\mathrm{tot}} = 0. \tag{27}$$

### ISOTHERMAL TITRATION CALORIMETRY (ITC)

A heat equation, based on Freiburger co-workers (2015) (4), can be calculated for each titration point:

$$\Delta H_j = \frac{(Q_j - Q_{j-1} + Q_{\mathrm{o}})}{V_{inj}[L]_{\mathrm{o}}} + \frac{(Q_j + Q_{j-1})}{2V_c[L]_{\mathrm{o}}}, \tag{28}$$

with $Q_j$ is the total energy as heat absorbed/released in the $j$-injection of ligand, $[L]_0$ the molar concentration of ligand added at each injection, $V_{inj}$ the volume of ligand injected, $V_c$ the working volume of the calorimeter cell, and $Q_{\mathrm{o}}$ is an offset due to the heat of mixing and dilution. The value of $Q_j$ is given by

$$Q_j = V_c[P]_{\mathrm{tot},j}\langle\Delta H_{bind}\rangle_j, \tag{29}$$

where $[P]_{\mathrm{tot},j}$ is the total protein concentration at each titration point, and $\langle\Delta H_{bind}\rangle_j$ is the molar enthalpy of binding. The latter can be computed as a weighted average of the enthalpy for the formation of $i$-state, $\Delta H_i$, using the probability weight defined in Eq. 3,

$$\langle\Delta H_{bind}\rangle_j = \sum_{i=0}^{n} \Delta H_i\, f_{PL_{i,j}}. \tag{30}$$

One should note that in this work it was assumed a mean molar binding enthalpy for every binding site, $\Delta H_{bind}$, such that the overall enthalpy for each $i$-liganded state is equal to $\Delta H_i = i\Delta H_{bind}$:

$$\langle\Delta H_{bind}\rangle_j = \sum_{i=0}^{n} i\Delta H_{bind}\, f_{PL_{i,j}} = \nu_j\Delta H_{bind}, \tag{31}$$

where $\nu_j$ is the average number of ligands bound per protein molecule at $j-$injection point, Eq. 7, calculated using the binding polynomial formalism. The assumption of a mean molar binding enthalpy is justified here because the four $Ca^{2+}$ binding sites are located at approximately 4Å from each other in the CALX-CBD1 crystallographic structure (5) and each $Ca^{2+}$ ion is coordinated by two or more electron donor groups from different amino acids.

### BAYESIAN STATISTICS

In the Bayesian approach, the probability represents a degree-of-belief of an assumption instead that of the Frequentist interpretation of the probability, which is related to the frequency with which an event occurs. Bayesian statistics is based in the Bayes' theorem: Let $\{d_i\}$ and $\mathbf{\Theta}(p_k)$ be the observed data and the theoretical model, respectively, where $\{p_k\}$ is the parameter set of the model. Thus, the probability that the model is correct given an experimental data set, i.e. the *posterior* is given by:

$$\mathcal{P}(\mathbf{\Theta}(p_k)|\{d_i\}, I) \propto \mathcal{L}(\{d_i\}|\mathbf{\Theta}(p_k), I)P(\mathbf{\Theta}(p_k)|I). \tag{32}$$

The term $P(\mathbf{\Theta}(p_k)|I)$ is called *prior* probability, it contains the initial available information about the model, and the term $\mathcal{L}(\{d_i\}|\mathbf{\Theta}(p_k), I)$ is called the *likelihood* function. The latter modifies the prior probability according to the data.

By using this approach one may calculate the marginalized probability distribution function of each parameter involved in the model. Therefore, the model's parameters are determined with statistical robustness (6, 7). One may also use prior information obtained from other experiments in the current investigation, e.g. results from calorimetry experiments could be used as prior input in the NMR analysis.

#### Fitting methodology

Fittings were performed using the `differential_evolution` function of the python package `scipy` to find the best fit and the initial position for Markov Chain Monte Carlo (MCMC). Samplings of the posteriori distribution functions of the fitted parameters were carried out through MCMC using the python package `emcee` (8). The reported parameter values correspond to the median of the marginalized distribution functions (PDF) sampled through MCMC. Reported uncertainties of the fitted parameters were taken from $1\sigma$ confidence interval (68%), centered at the percentile 50% of the cumulative distribution function (CDF). In order to ensure convergence of the chains we used the following conditions for each analysis:

1. ITC: 100 walkers, 4000 (Hill) - 5000 (stepwise) points and 3000 (Hill) - 4000 (stepwise) of burn-in.
2. NMR titration: 100 walkers, 1000 points and 600 of burn-in.
3. CPMG: 80 walkers, 1300 points and 800 of burn-in.
4. CEST: 100 walkers, 1200 points and 600 of burn-in.

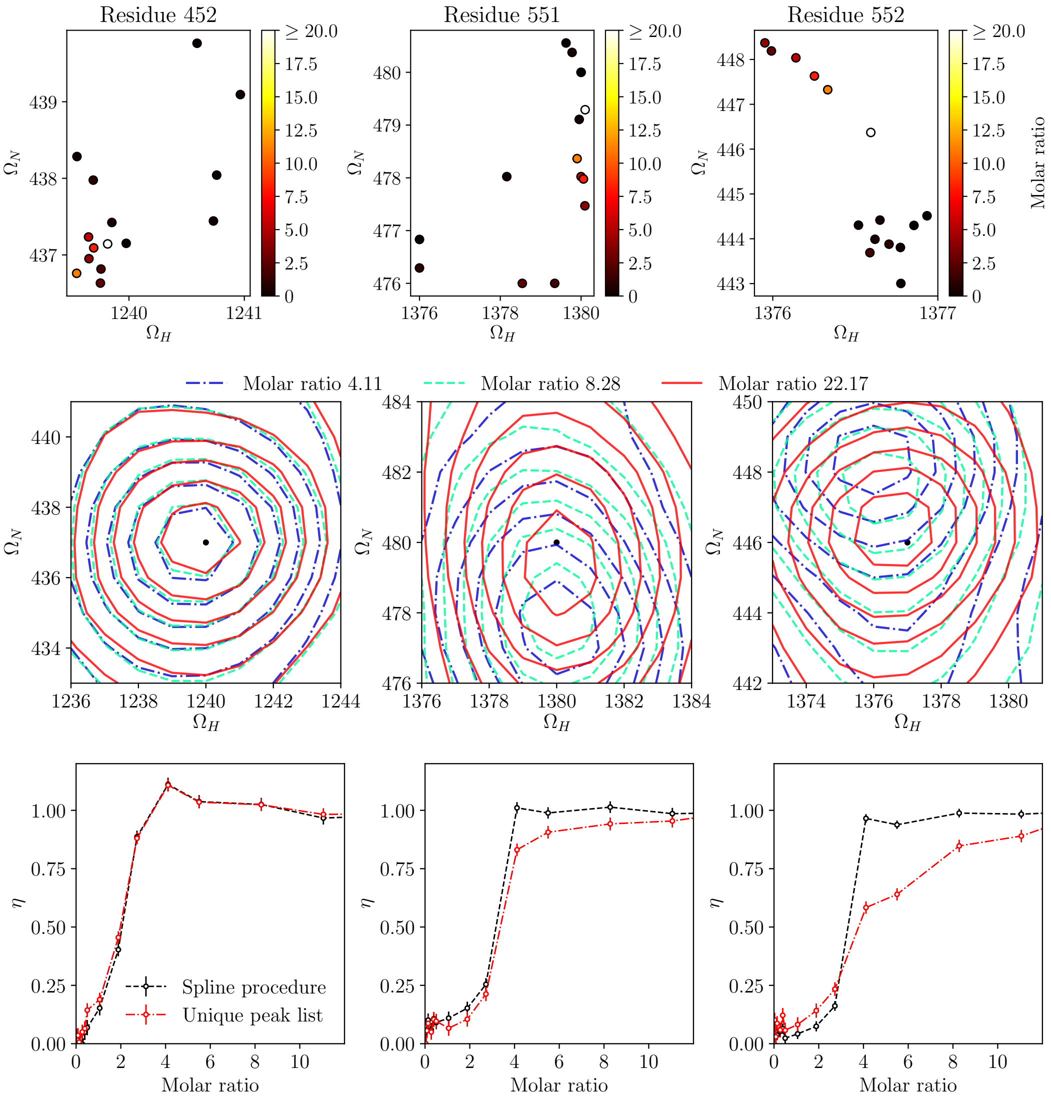


Supplementary Figure 1: Representative examples of the spline procedure adopted to determine the HSQC cross peak intensities in the NMR titration experiment. The examples shown are for residues 452 (left), 551 (middle) and 552 (right). Top: scatter plots of the center of the peaks determined by spline as a function of the $Ca^{2+}$ concentration. Middle: level curves for the two-dimensional peaks with normalized intensities at different $Ca^{2+}$ concentrations. The black dots indicate the peak position at $Ca^{2+}$-saturating concentrations. Bottom: $Ca^{2+}$-binding curves determined assuming that the peak position does not move with respect to its final position at saturating $Ca^{2+}$-concentrations (red), or assuming that it moves and determining the intensity using *spline* (black). All plots are shown as matrix position instead of frequency units. Clearly, unlike for residue 452, very different $Ca^{2+}$-binding curves would be obtained for residues 551 and 552 in case the cross peak position was assumed to be fixed at the position found at saturating $Ca^{2+}$-concentrations.

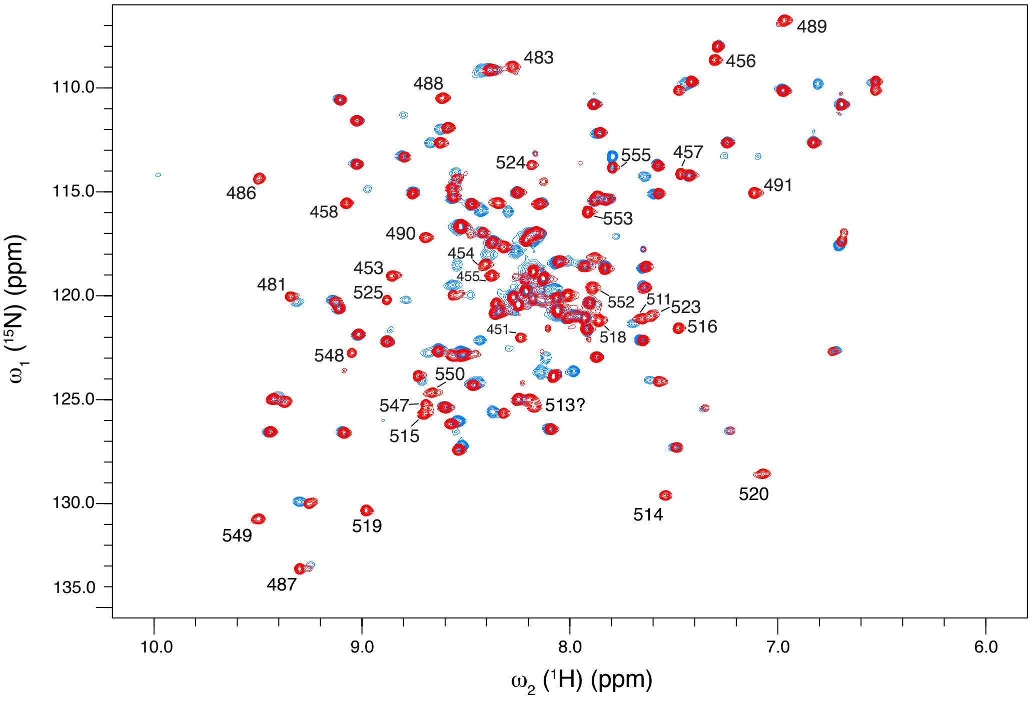


Supplementary Figure 2: Superposition of $^{1}$H-$^{15}$N HSQC spectra of CALX-CBD1 recorded in the absence (blue) and presence (red) of 20 mM of $CaCl_2$. Assignments for the peaks that appear only in the presence of $Ca^{2+}$ are indicated. The NMR spectra were recorded at 800 MHz ($^{1}$H frequency) and 308 K, as matrices of 768x128 complex points. Protein samples consisted of approximately 200 $\mu$M of $^{15}$N labeled CALX-CBD1. The Apo state sample contained 2 mM of EDTA, while the $Ca^{2+}$-bound sample contained 20 mM of $CaCl_2$.

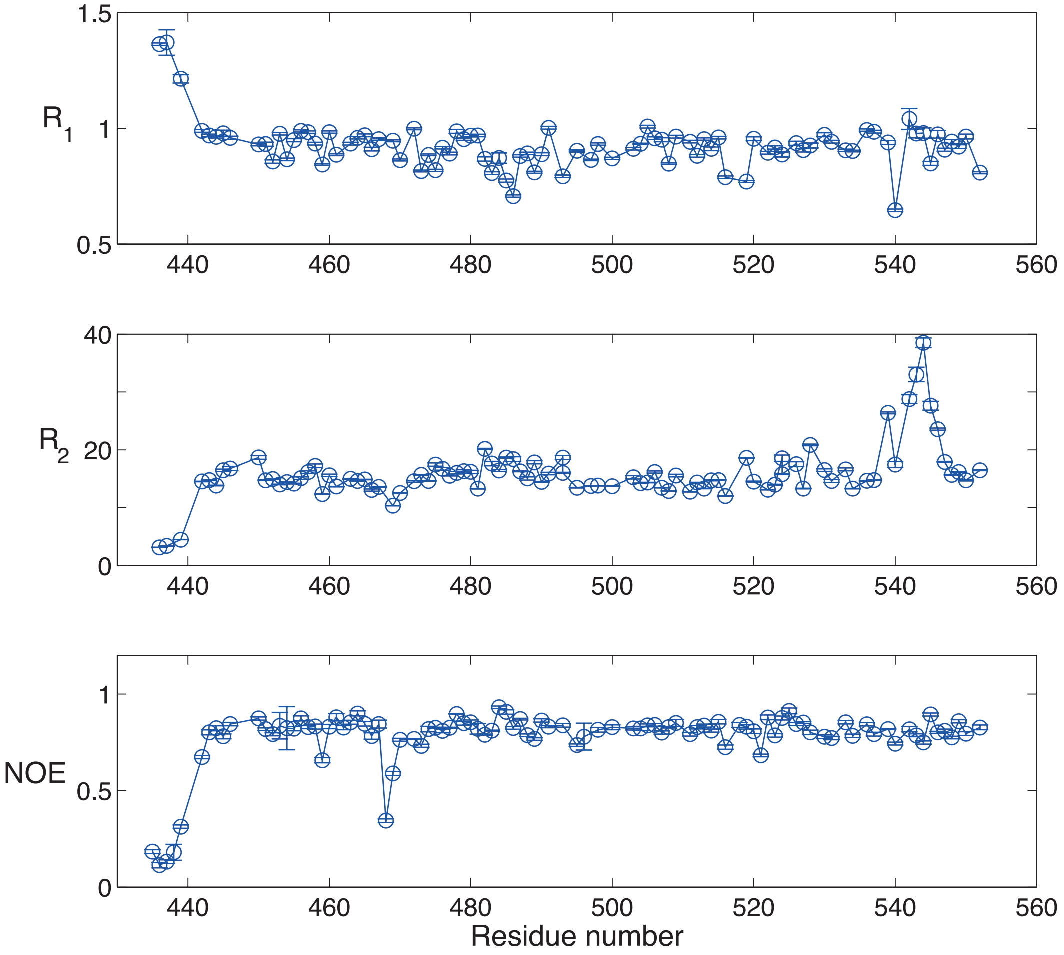


Supplementary Figure 3: $^{15}N$ transverse ($R_2$) and longitudinal ($R_1$) magnetization relaxation rates, and the heteronuclear NOE as a function of the amino acid sequence. The protein sample consisted of 380 $\mu$M of $^{15}N$-labeled CALX-CBD1 containing $CaCl_2$ at a molar ratio of 16:1 ($CaCl_2$:protein) (Holo state). The $^{15}N$ relaxation data were recorded at 800 MHz ($^1H$ frequency) and 308 K. Data acquisition, analysis, and error propagation are explained in the "Methods section" in the main text.

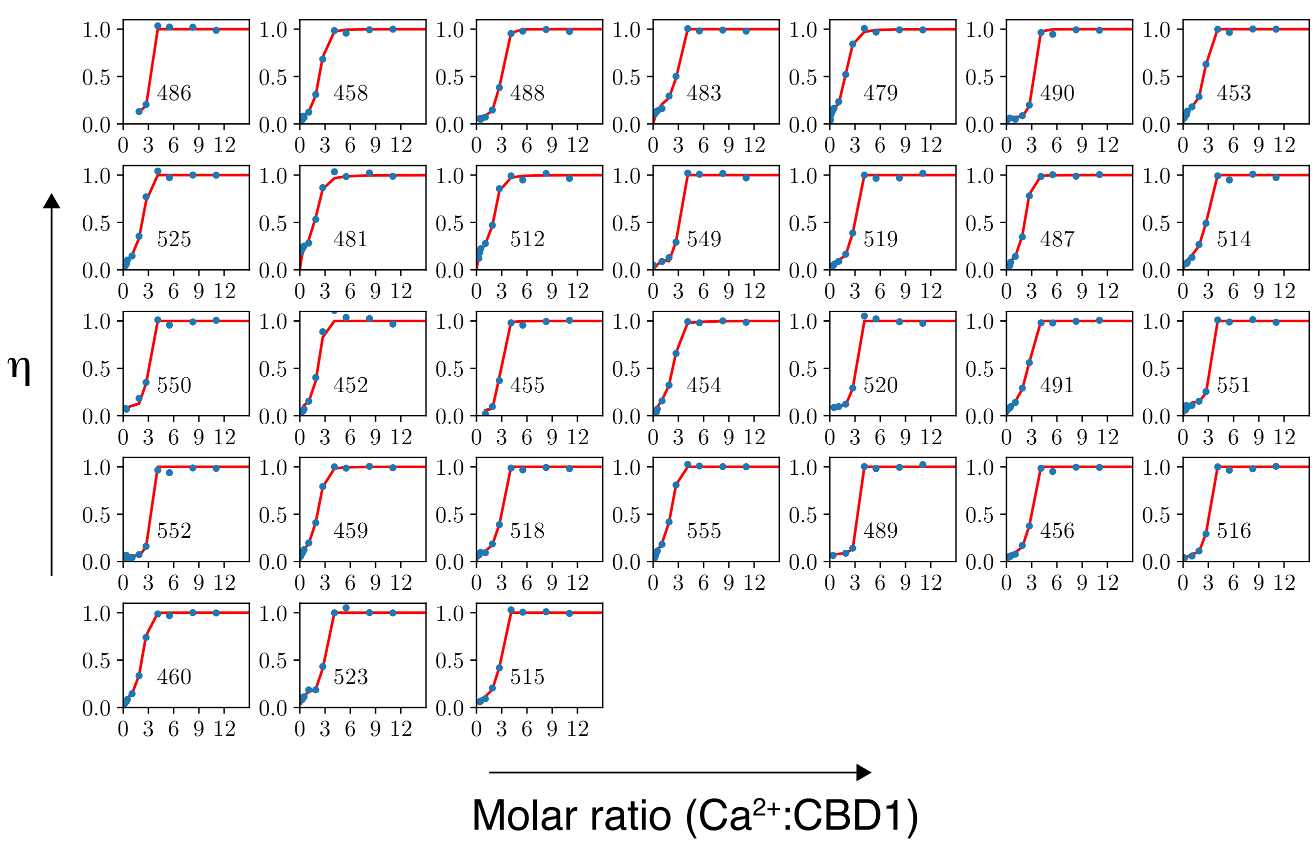


Supplementary Figure 4: NMR binding curves obtained for selected residues and calculated according to Eq. 1 (Main text) (blue), and fittings to Eq. 7 (Main text) (red). The residue number is indicated inside of each plot.

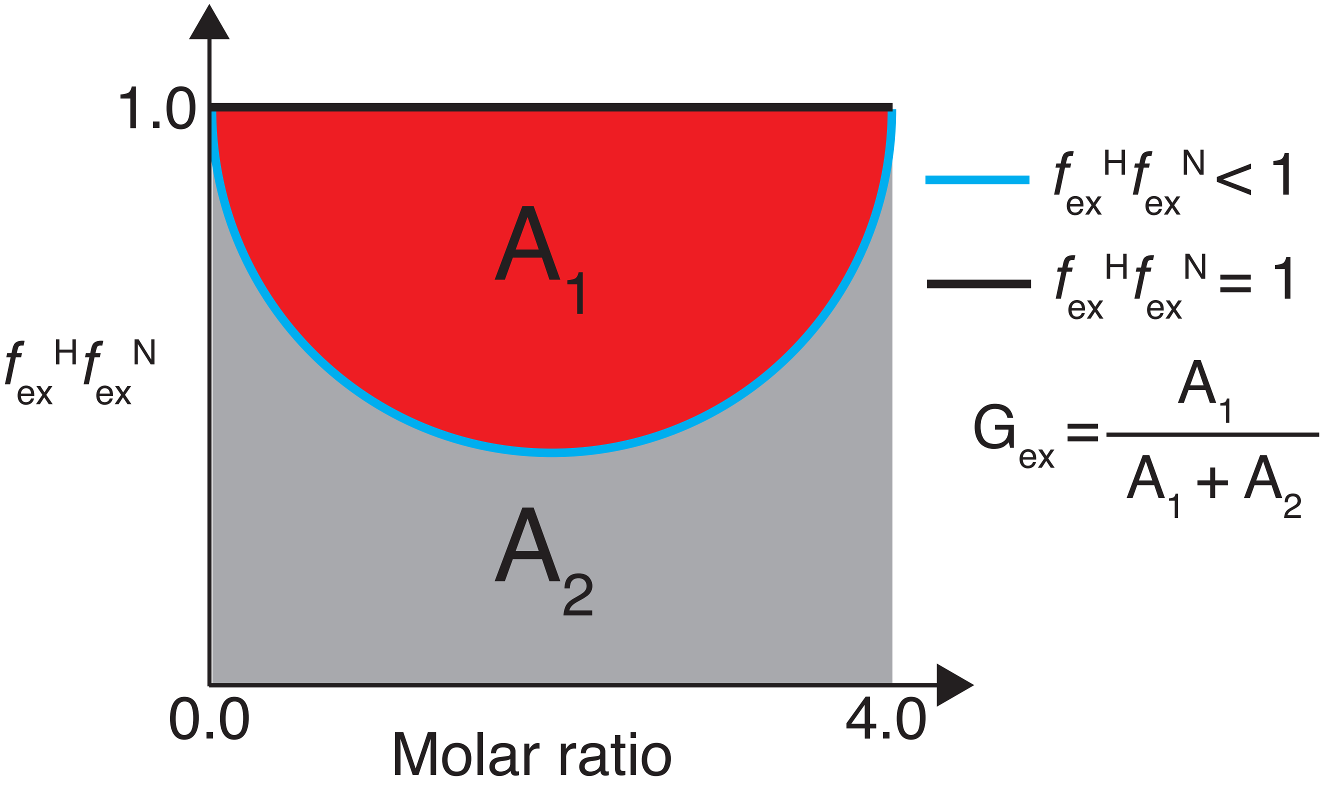


Supplementary Figure 5: The factor $G_{ex}$ is a geometrical interpretation of the magnitude of the chemical exchange contributions to the binding curve. $G_{ex}$ is calculated as the normalized $A_1$ area. $A_2$ is defined as the area under the $f_{ex}^{H} f_{ex}^{N}$ function (blue line), while $A_1 + A_2$ is the area under the rectangle (black line) calculated from molar ratios 0 to 4.

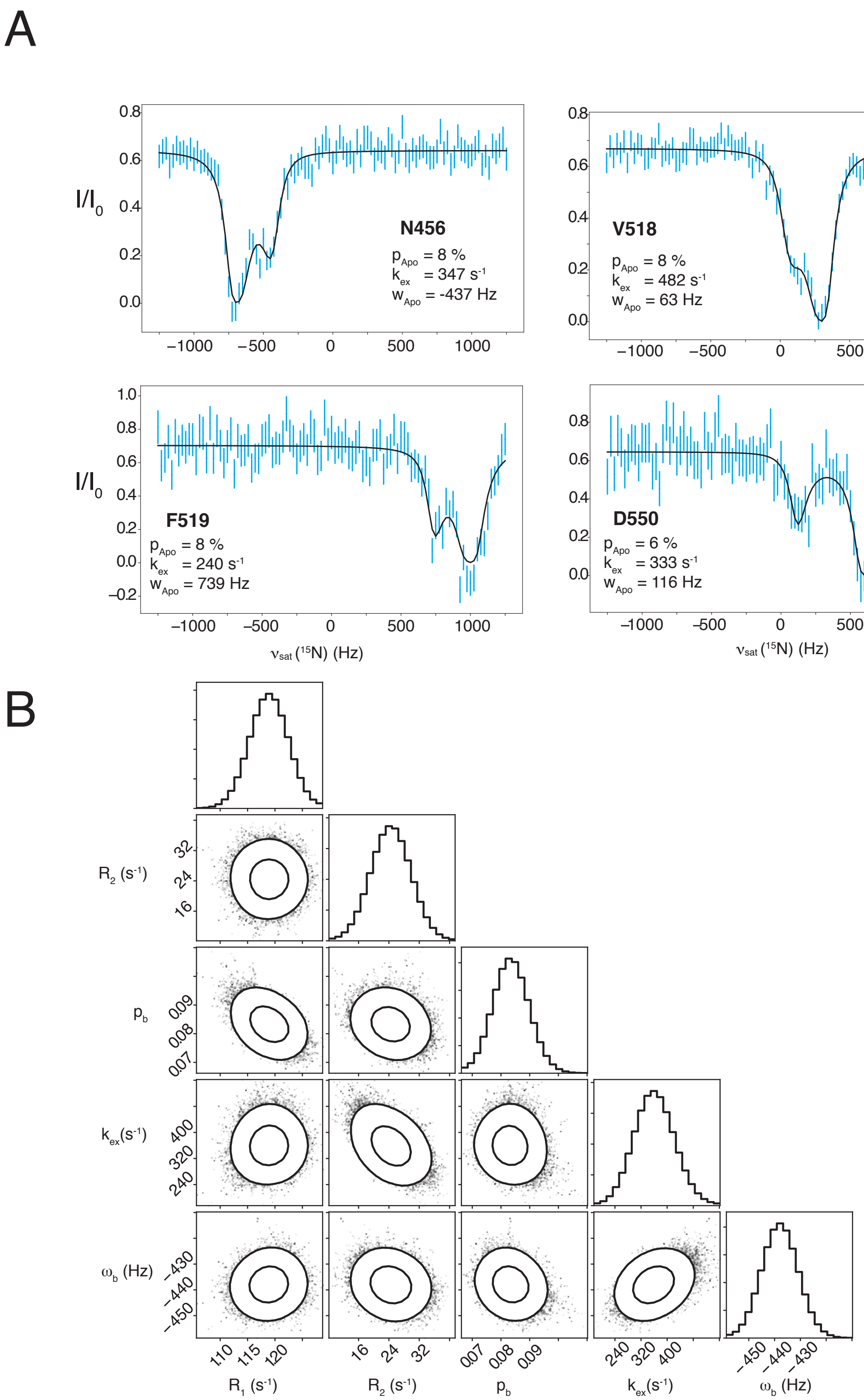


Supplementary Figure 6: **(A)** $^{15}$N-CEST profiles obtained for CALX-CBD1 in the presence of $CaCl_2$ at the molar ratio of 4:1 ($Ca^{2+}$:CALX-CBD1) (cyan), superimposed on the fittings to the Bloch-McConnell equation (black). The uncertainties and the experimental conditions are explained in the main text. **(B)** Example of one dimensional maginalized parameter distributions and two-dimensional correlation plots, obtained by fitting the CEST profile of Asn456 to the Bloch-McConnell equation.

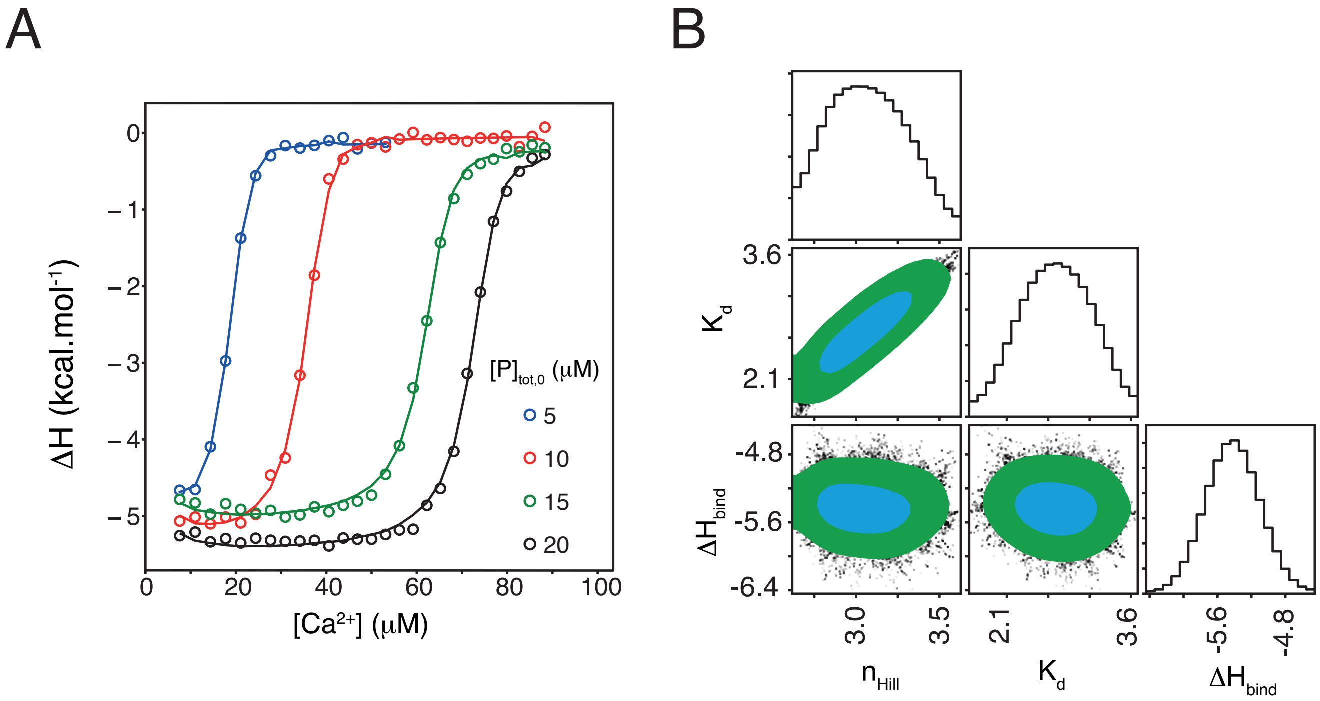


Supplementary Figure 7: A) Calorimetric titrations of CALX-CBD1 with $CaCl_2$ using different protein concentrations. The lines represent fittings to the Hill model. B) One dimensional marginalized parameter distributions and 2D marginalized distributions of pairs of parameters. The $1\sigma$ confidence contour region is shown in cyan, while the $2\sigma$ confidence contour region is shown in green plus cyan.

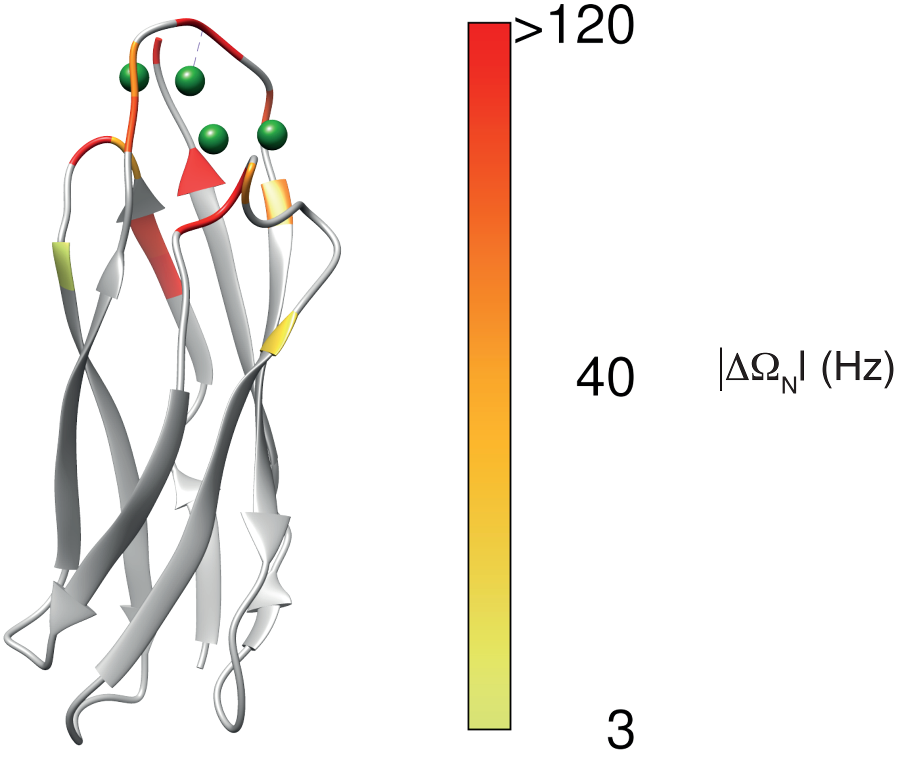


Supplementary Figure 8: Chemical shift differences ($\Delta\Omega_N$) between CALX-CBD1 in the Apo and in the fully $Ca^{2+}$-bound state were mapped on the crystal structure of CALX-CBD1 (PDB 3E9T). $Ca^{2+}$ ions are shown as green spheres. $\Delta\Omega_N$ was obtained by fitting the CPMG relaxation dispersion curves to the Bloch-McConnell equation assuming a two-site exchange model. The relaxation dispersion experiments were recorded at two static magnetic fields corresponding to 800 and 600 MHz ($^1$H frequency), on a sample of CALX-CBD1 in the presence of $CaCl_2$ at the 1:4 molar ratio (CALX-CBD1:$CaCl_2$). The protein concentration was 380 $\mu$M.
